# Internal state dependent control of feeding behaviour via hippocampal ghrelin signalling

**DOI:** 10.1101/2021.11.05.467326

**Authors:** Ryan W. S. Wee, Karyna Mishchanchuk, Rawan AlSubaie, Andrew F. MacAskill

**Affiliations:** Department of Neuroscience, Physiology and Pharmacology, University College London, Gower St, London, WC1E 6BT

## Abstract

Hunger is an internal state that not only invigorates feeding, but also acts as a contextual cue for the higher-order control of anticipatory feeding-related behaviour. The ventral hippocampus is a brain region crucial for differentiating optimal behaviour across different contexts, but how internal context such as hunger influence hippocampal circuits to define behaviour is not known. Pyramidal neurons in the ventral hippocampus, including the ventral CA1/subiculum border (vS) express the receptor for the peripheral hunger hormone ghrelin, and ghrelin is known to cross the blood brain barrier and directly influence hippocampal circuitry. But how ghrelin influences vS has not been directly investigated. In this study, we used a combination of electrophysiology, optogenetics and *in vivo* calcium imaging in mice to investigate the role of vS during feeding behaviour across different states of hunger. We found that activity of a unique subpopulation of vS neurons that project to the nucleus accumbens (vS-NAc) increased when animals approached and investigated food, and this activity inhibited the transition to begin eating. Increases in peripheral ghrelin reduced vS-NAc activity during this anticipatory phase of feeding behaviour by increasing the postsynaptic influence of inhibition, and promoted the initiation of eating. Furthermore, this peripheral ghrelin-induced inhibition required postsynaptic expression of the ghrelin receptor GHSR1a in vS-NAc neurons, and removal of GHSR1a from vS-NAc neurons impaired ghrelin-induced changes in feeding-related behaviour. Together, these experiments define a ghrelin-sensitive hippocampal circuit that informs the decision to eat based on internal state.

## INTRODUCTION

Animals must be able to control feeding behaviour dependent on need. Consuming food when already sated utilises time and energy that could be spent on more essential functions, and can result in disease and disorders associated with overeating. In contrast, being unable to sense the need for food – or ‘hunger’ - can result in undereating, and resultant lack of fitness (Toates, 1981).

A key aspect of this process is the ability to integrate external cues with internal state such as hunger (Benoit et al., 2010; Burnett et al., 2019; Toates, 1981). This is because the value of a given food cue is ambiguous - the same food would predict a rewarding post ingestive outcome when the animal is hungry, but not when the animal is sated (Davidson et al., 2007). In this framework, hunger can act as a context upon which the optimal behaviour towards sensory cues is interpreted (Azevedo et al., 2019; Davidson and Jarrard, 1993; Davidson et al., 2007; Gershman, 2017; Mohammad et al., 2021; Rudy and Sutherland, 1995).

The hippocampus, and particularly its output area the ventral CA1 / subiculum (vS) – has been repeatedly proposed as a crucial structure for defining behaviour dependent on context, most notably in spatial contextual associations (Good and Honey, 1991; Holland and Bouton, 1999; Per Andersen et al., 2006; Strange et al., 2014). However, the hippocampus is also heavily involved in hunger sensing, in both humans and rodents (Azevedo et al., 2019; Davidson and Jarrard, 1993; Davidson et al., 2007; Mohammad et al., 2021; Wallner-Liebmann et al., 2010; Wang et al., 2006). This suggests that in addition to spatial context, the hippocampus may also differentiate behaviour dependent on other, more abstract contexts such as hunger. Consistent with this idea, hippocampal activity is extremely sensitive to hunger state in both humans (Wallner-Liebmann et al., 2010; Wang et al., 2006) and rodents (Carey et al., 2019; Kennedy and Shapiro, 2009; Min et al., 2011), and inactivation and dysfunction of the hippocampus leads to impaired hunger-based decision making (Azevedo et al., 2019; Davidson and Jarrard, 1993; Hebben et al., 1985; Mohammad et al., 2021; Rozin et al., 1998; Scoville and Milner, 1957).

In keeping with the hippocampus’ role in responding to internal state, hippocampal pyramidal neurons express a diverse array of physiologically important receptors, for example, those involved in the signalling axes for stress, hunger and thirst (Lathe, 2001). More specifically to hunger, the hippocampus expresses the receptor for the peripheral hunger hormone ghrelin (GHSR1a) in both rodents (Diano et al., 2006; Guan et al., 1997; Hsu et al., 2015; Mani et al., 2014; Zigman et al., 2006) and non-human primates (Mitchell et al., 2001). Interestingly, many peripherally circulating hormones are able to gain access to the hippocampus (Hamasaki et al., 2020), and there is evidence to support the entry of peripheral ghrelin into the hippocampus through the blood-brain barrier (Banks et al., 2002; Diano et al., 2006, but see Furness et al., 2011). Once present in the hippocampus, ghrelin is capable of not only inducing structural and functional plasticity (Diano et al., 2006; Ribeiro et al., 2014), but also influencing anticipatory behaviour and influencing choice (Diano et al., 2006; Hsu et al., 2015, 2016; Kanoski et al., 2013, Yang et al., 2020).

However, while it is clear that motivational state affects hippocampal processing, and hippocampal dysfunction impairs hunger dependent behaviour, how hippocampal circuitry directly influences internal state dependent feeding behaviour, and the cellular and circuit mechanisms underlying this ability remains unknown. This is compounded by the fact that vS is composed of multiple, non-overlapping and functionally distinct parallel projections to distinct downstream areas (AlSubaie et al., 2021; Cembrowski and Spruston, 2019; Gergues et al., 2020; Sánchez-Bellot et al., 2022; Soltesz and Losonczy, 2018; Wee and MacAskill, 2020). For example, neurons in vS that project to the nucleus accumbens (NAc) have been shown to preferentially represent and control motivation and value (AlSubaie et al., 2021; Britt et al., 2012; LeGates et al., 2018; Trouche et al., 2019), to be preferentially active during anticipatory behaviour (Ciocchi et al., 2015), and inhibited upon eating (Reed et al., 2018). Similarly, a separate population of neurons projecting to the lateral hypothalamus (LH) has been shown to be recruited during salient environments and during learning of food associations (Hsu et al., 2015; Jimenez et al., 2018; Mohammad et al., 2021). Both of these populations of neurons are therefore well placed to control anticipatory feeding related behaviour (AlSubaie et al., 2021; Britt et al., 2012; Hsu et al., 2015; Jimenez et al., 2018; LeGates et al., 2018; Mohammad et al., 2021; Reed et al., 2018; Trouche et al., 2019). However, how these populations are uniquely used during feeding behaviour, and how they are influenced by internal state signalled by peripheral ghrelin is unknown.

Together, vS is well placed to control anticipatory feeding-related behaviour. It is consistently implicated in hunger-based decisions, its dysfunction impairs behaviour requiring hunger sensing, and it has ghrelin-sensitive neurons that project to two brain regions both crucially important for defining feeding behaviour. Therefore, in this study we used a combination of quantitative behaviour, *in vivo* imaging and manipulation, and slice physiology to address the role of vS circuitry in hunger-based decisions. We found that activity of a unique subpopulation of vS neurons that project to the nucleus accumbens (vS-NAc) increased when animals approached and investigated food, and this activity inhibited the transition to begin eating. Increases in peripheral ghrelin reduced vS-NAc activity during this anticipatory phase of feeding behaviour by increasing the postsynaptic influence of inhibition. Furthermore, removal of the ghrelin receptor (GHSR1a) specifically from vS-NAc neurons impaired ghrelin-induced changes in feeding-related behaviour. Together, these experiments define a ghrelin-sensitive hippocampal circuit that informs the decision to eat based on internal state.

## RESULTS

### Peripheral ghrelin administration increases the transition from food investigation to food consumption

Feeding behaviour can be described as the chaining together of distinct, stereotyped behaviours such as exploratory sniffing and investigation of food (presented here as ‘Inv’) food consumption (‘Eat’), as well as non-feeding behaviours such as ‘Rear’, ‘Groom’ and ‘Rest’ (‘Oth’) (Halford et al., 1998). Increases in peripheral ghrelin are known to dramatically alter behaviour towards food, in particular, through the promotion of the initiation of eating. However, despite intensive investigation, it is unclear how increases in peripheral ghrelin alter the structure of this moment-to-moment behaviour around food.

To address this, we first confirmed that ghrelin injections caused an increase in food consumption in sated mice when they were repeatedly presented with a familiar food item (a chow pellet) in a well habituated arena, when compared to injection of PBS vehicle control (**Figure 1A**). Next, by scoring each behaviour performed by the mouse during the session, we found that this increased consumption resulted from a large and specific increase in the frequency of initiating eating, with only minimal change in the frequency of food investigation or the frequency of non-feeding behaviours (**Figure 1B**).

**Figure 1.**
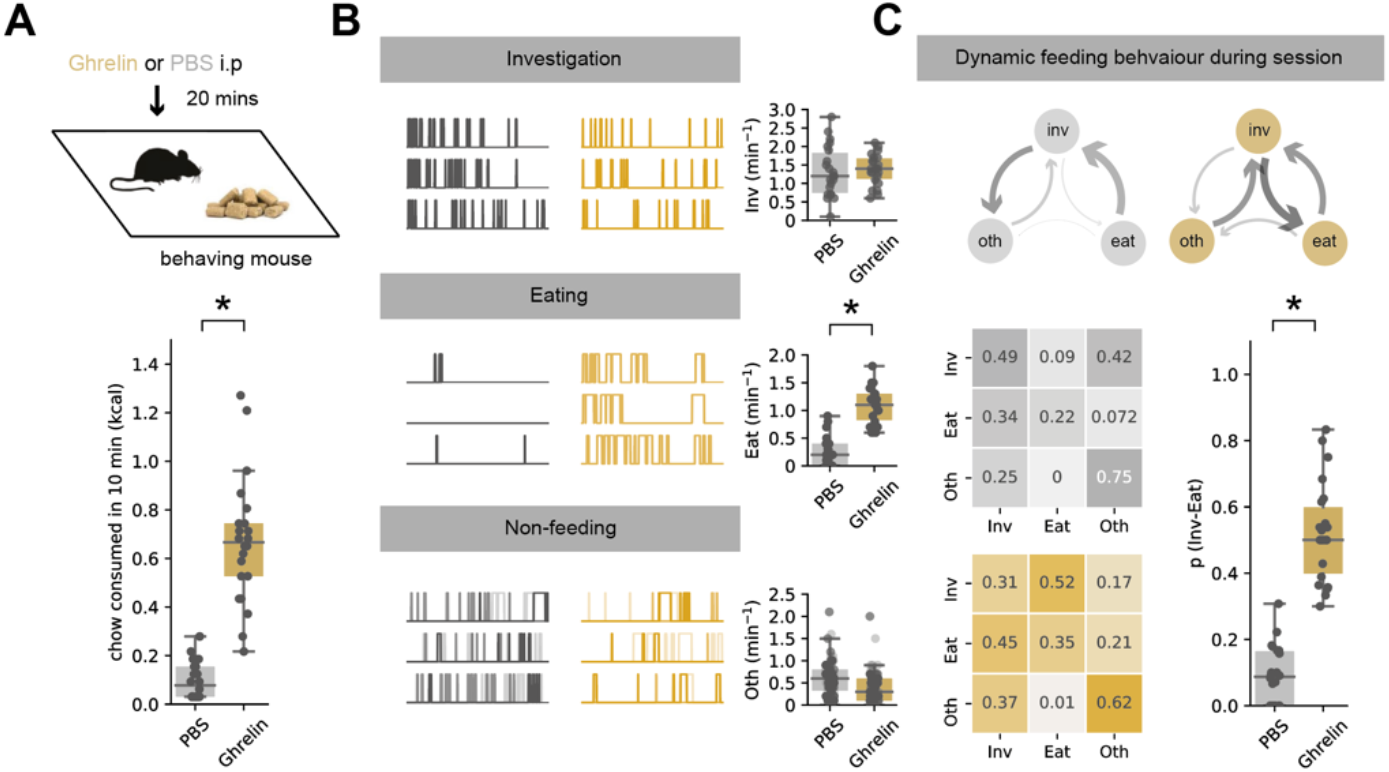
Peripheral ghrelin administration increases the transition from food investigation to food consumption. **A**.Top, schematic of experiment. Mouse was injected with either ghrelin or control PBS before exploring a well habituated chamber containing familiar chow for 10 min. Bottom, ghrelin administration (gold) increases chow consumption compared to PBS control (grey). **B**. Analysis of food investigation, eating, and non-feeding behaviours including grooming, quiet resting and rearing. Plots show three examples of mouse behaviour across example ten-minute sessions. Note that overall ghrelin has minimal effect on behaviour apart from large increases in number of eating bouts. **C**. Markov analysis of feeding behaviour during ten-minute session. Top, state transitions for PBS (grey) and ghrelin (gold) treated mice. Arrow thickness is proportional to the probability of transition. Bottom, left, transition matrix for PBS and ghrelin treated mice. Each column and row add to one, and so the matrix represents the probability of the next behavioural state given the current state. Note that ghrelin increases the transition from investigation to eating, with minimal influence on other behavioural transitions. Bottom, right, summary of investigation to eat transition across all mice in PBS and ghrelin. Note large increase after ghrelin in all mice.

Next, we asked how sequences of behaviour changed within each session to result in this increase in eating. To quantitatively describe the organisation of such sequences, we analysed scored behaviour (‘Inv’, ‘Eat’ and ‘Oth’) as a discrete-time Markov chain - a vector of behavioural states that change as a function of time (Burnett et al., 2019). We then computed the transition matrix P*ij* for each animal (which defines the animal’s probability of transitioning from behaviour *i* to behaviour *j* during the session), and compared these matrices across different states of hunger (**Figure 1C**).

Using this analysis, we found that the effect of ghrelin was very specific, and was centred around transitions from food investigation (‘Inv’). While PBS-treated mice frequently investigated the food pellet, this investigation was very rarely followed by a transition to eating. In contrast, in ghrelin treated mice, the frequency of investigation of the food pellet was not changed (**Figure 1B, Supp Fig 1**), but the transition from investigation to eating was substantially increased. Overall, this suggests that the effect of ghrelin was to increase the probability of transitioning from investigation of, to consumption of food (p(Inv->eat)), with only minimal effect on other behavioural transitions (**Figure 1C, Supp Fig 2**). This specific alteration in behaviour can account for previous work suggesting a key role of ghrelin signalling in the anticipation and initiation of feeding.

**Figure 2.**
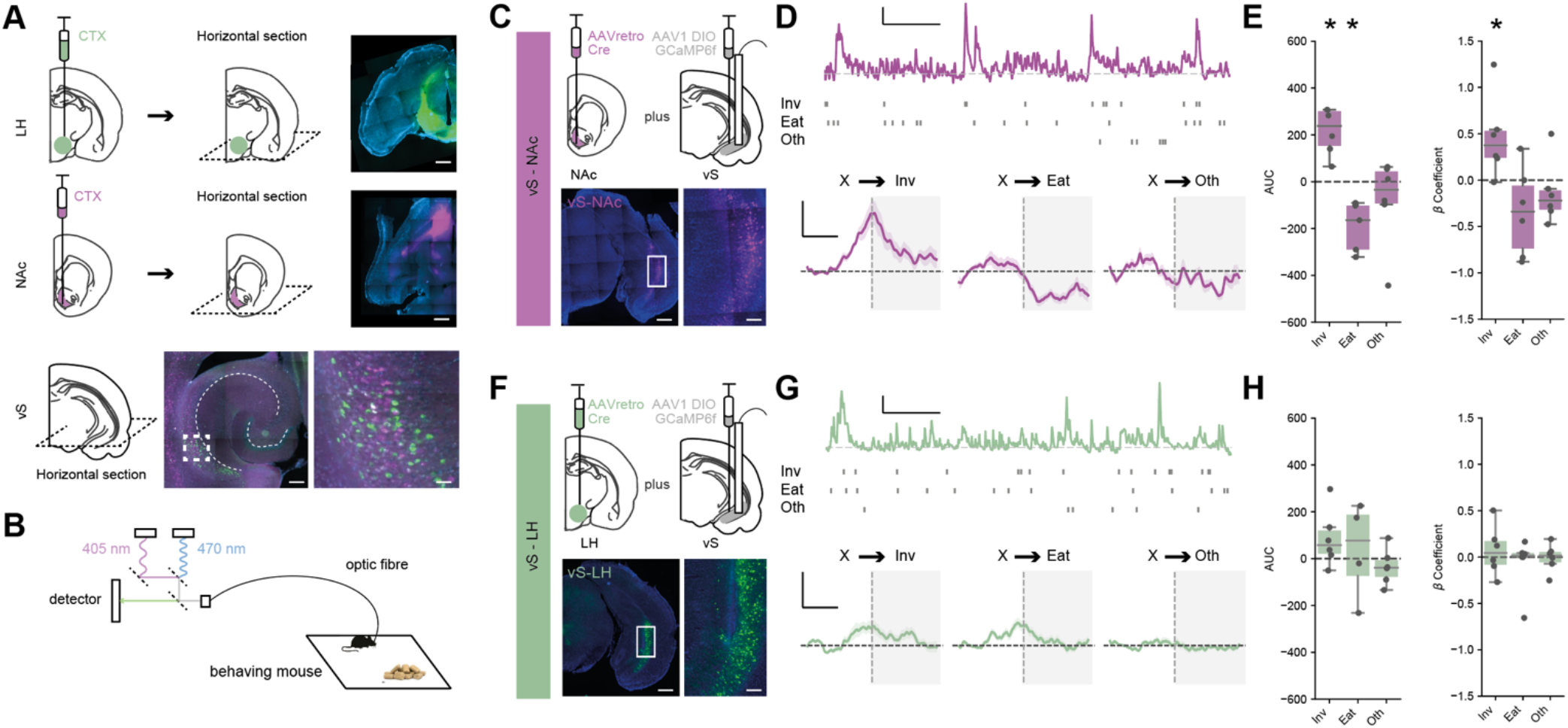
vS neurons that project to NAc are active during investigation of food. **A** Retrograde labelling of vS projections to LH and NAc identified by injections of cholera toxin (CTX). Top, injection in LH (green) and NAc (purple) shown in horizontal slices 2 weeks after injection. Bottom, retrogradely labelled neurons in vS. Note two populations with only minimal overlap. Scale bars: Top = 500 µm, Bottom, 200 µm, 25 µm zoom. **B**. Schematic of photometry setup to allow bulk imaging during free behaviour. **C**. Top, schematic of injections allowing intersectional targeting of vS-NAc neurons for photometry. Bottom example images of neurons in vS. Scale bars = 500 µm, 100 µm zoom. **D**. Top, example photometry trace from vS-NAc neurons during the session, with start point of Investigative (Inv), Eating (Eat) and Non feeding (Oth) behavioural events plotted as raster plots below. Scale bar = 1 zF, 2 min. Bottom, average activity for vS-NAc neurons across all mice aligned to start of behavioural events during the session. Note large increase in activity around investigative bout and decrease in activity on initiation of eating. Scale bar = 0.5 zF, 4 s. **E**. Summary of activity around each behaviour for vS-NAc neurons, summarised using either the area under the curve (AUC) of event-aligned activity (left), or using the coefficients of a generalised linear model fir to the calcium data (right, see methods). Note consistent increase in activity around investigation in vS-NAc neurons using both metrics. **F-H**. As in **D-E** but for vS-LH neurons. Note lack of consistent activity aligned to behavioural events.

### vS-NAc activity tracks investigation and eating behaviour

Despite much investigation, the precise neural mechanisms underlying the effect of ghrelin on anticipatory feeding behaviour remain unclear. This problem is often characterised as one of conditional ambiguity (Davidson et al., 2007; Gershman, 2017) – where the appropriate behaviour towards the same food-associated cue is different dependent on internal state. One area that is thought to play a key role in this ability is the ventral hippocampus, via its strong excitatory projections from the CA1/subiculum border (vS) to downstream areas such as the nucleus accumbens (NAc) (AlSubaie et al., 2021; Britt et al., 2012; LeGates et al., 2018; Trouche et al., 2019; Wee and MacAskill, 2020), and the lateral hypothalamus (LH) (Hsu et al., 2015; Jimenez et al., 2018; Mohammad et al., 2021; Wee and MacAskill, 2020) that arise from distinct, minimally overlapping population of neurons (Gergues et al., 2020; Naber and Witter, 1998; Wee and MacAskill, 2020). Therefore, we next investigated i) how activity in vS was modulated during feeding behaviour, and ii) how this activity was distributed across the two populations of projection neurons.

We first confirmed that the two projections from vS were composed of largely non overlapping populations using retrograde cholera toxin labelling (**Figure 2A**, Wee and MacAskill, 2020).

Next, we used intersectional viral expression of cre-dependent GCaMP6f in vS, paired with an injection of AAVretro-cre in either NAc or LH, and combined this with implantation of an optical fibre immediately above vS (**Figure 2C,F**). This allowed us to record bulk calcium activity of each projection pathway as mice were freely behaving in response to presentation of chow (**Figure 2B**).

For each mouse, we first aligned this calcium activity to the onset of each class of behaviour (either ‘Inv’, ‘Eat’ or ‘Oth’, **Figure 2D**). We noticed that there was a large and consistent increase in the activity of vS-NAc neurons leading up to and during the investigation of food, reminiscent of previous descriptions of these neurons ramping towards salient locations (Ciocchi et al., 2015, **Figure 1D,E**). In addition, there was a consistent drop of vS-NAc activity upon the initiation of eating (Reed et al., 2018). Together this suggests that vS-NAc neurons are active during anticipation and investigation of food, but are then rapidly inhibited upon the commencement of feeding. In contrast, despite multiple suggestions of a crucial role for vS-LH neurons in learning about food dependent, and other affective cues, in this simple assay vS-LH neurons did not show any consistent activity that was time locked to exploratory or feeding behaviour in this task (**Figure 2G,H**). Instead vS-LH neurons showed robust activity in response to salient stimuli such as the presentation of an object or chow (González et al., 2016, **Supp Fig 3**).

**Figure 3.**
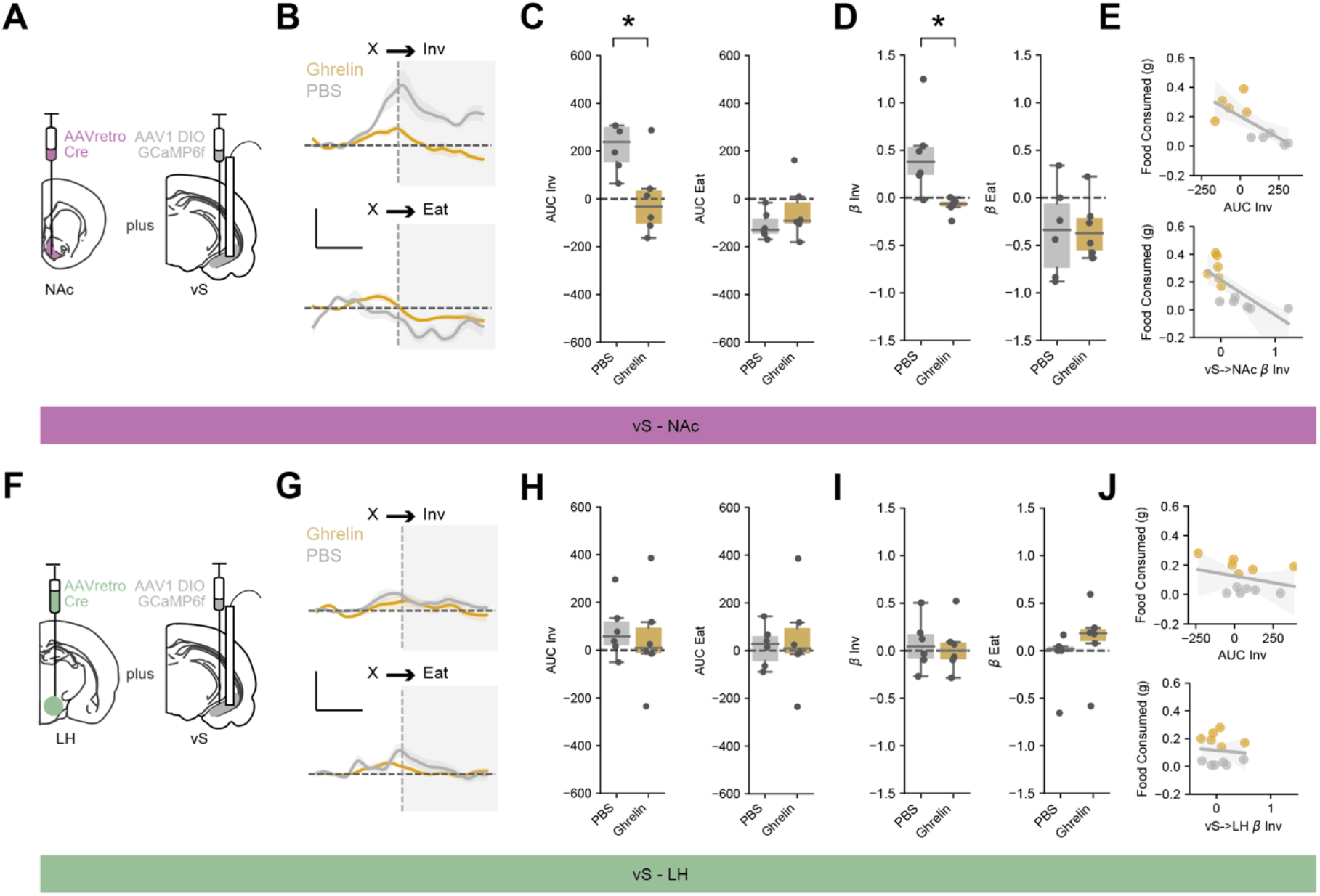
Increased peripheral ghrelin inhibits vS-NAc activity during food investigation. **A** Schematic of injections allowing intersectional targeting of vS-NAc neurons for photometry. **B**. Top, average activity for vS-NAc neurons across all mice aligned to investigation, after injection of either PBS (grey) or ghrelin (gold). Bottom, average activity around eating. Note large increase in activity around investigation after PBS, but not after ghrelin, and no change in activity around eating. Scale bar = 0.5 zF, 4 s. **C, D**. Summary of activity around investigation (left) and eating (right) for vS-NAc neurons after injection of either PBS (grey) or ghrelin (gold). **C** shows event-aligned AUC, **D** shows coefficients of a generalised liner model. In both analyses, note increase in activity after PBS around investigation is not present after ghrelin, while decrease in activity during eating is similar in both conditions. **E**. Correlation between vS-NAc activity during investigation and chow consumption using AUC (top) or coefficients (bottom). Note strong inverse relationship between vS-NAc activity and food consumption using both measures. **F-J**. As (**A-E**) but for vS-LH neurons. Note lack of activity around behavioural events that is also insensitive to ghrelin, and no correlation between vS-LH activity and food consumed in the session.

Together, this suggests that the activity of vS-NAc neurons is bidirectionally modulated during investigation and consumption of familiar food. Moreover, increases in vS-NAc activity around the investigation of food is consistent with theoretical descriptions of a role in the anticipation of feeding.

### vS-NAc activity during food investigation is inhibited by ghrelin

Our previous results revealed that there was a large anticipatory ramp of activity in vS-NAc neurons during investigation of food (**Figure 2**). As the effect of ghrelin was to alter the consequences of such investigative behaviour (**Figure 1**), we next wanted to investigate how this activity in vS was modulated by increases in peripheral ghrelin.

We repeated our investigation of vS activity in mice with counterbalanced injections of either ghrelin or PBS (**Figure 3**). As before, ghrelin injections markedly increased both total consumption of chow, but also specifically the transition from investigation to eating in both cohorts of mice (**Supp Fig 4**). Importantly however, this change in behaviour was accompanied by a substantial reduction of the activity of vS-NAc neurons during food investigation (**Figure 3B-D**). This effect was specific to activity around investigation, as alterations in vS-NAc activity upon eating were maintained across both ghrelin- and PBS-treated animals. We reasoned that the behavioural effect of ghrelin – increased chow consumption due to an increased transition from investigation to eating – may be defined by the magnitude of inhibition of vS-NAc neurons. Consistent with this idea, on a session-by-session basis there was a negative correlation between the activity of vS-NAc neurons during food investigation and the total amount of food consumed in that session (**Figure 3E**). Interestingly, again there was no effect of ghrelin on vS-LH neurons, and activity remained invariant across each behaviour (**Figure 3F-J**), suggesting a projection specific modulation of vS neurons upon increases of peripheral ghrelin.

**Figure 4.**
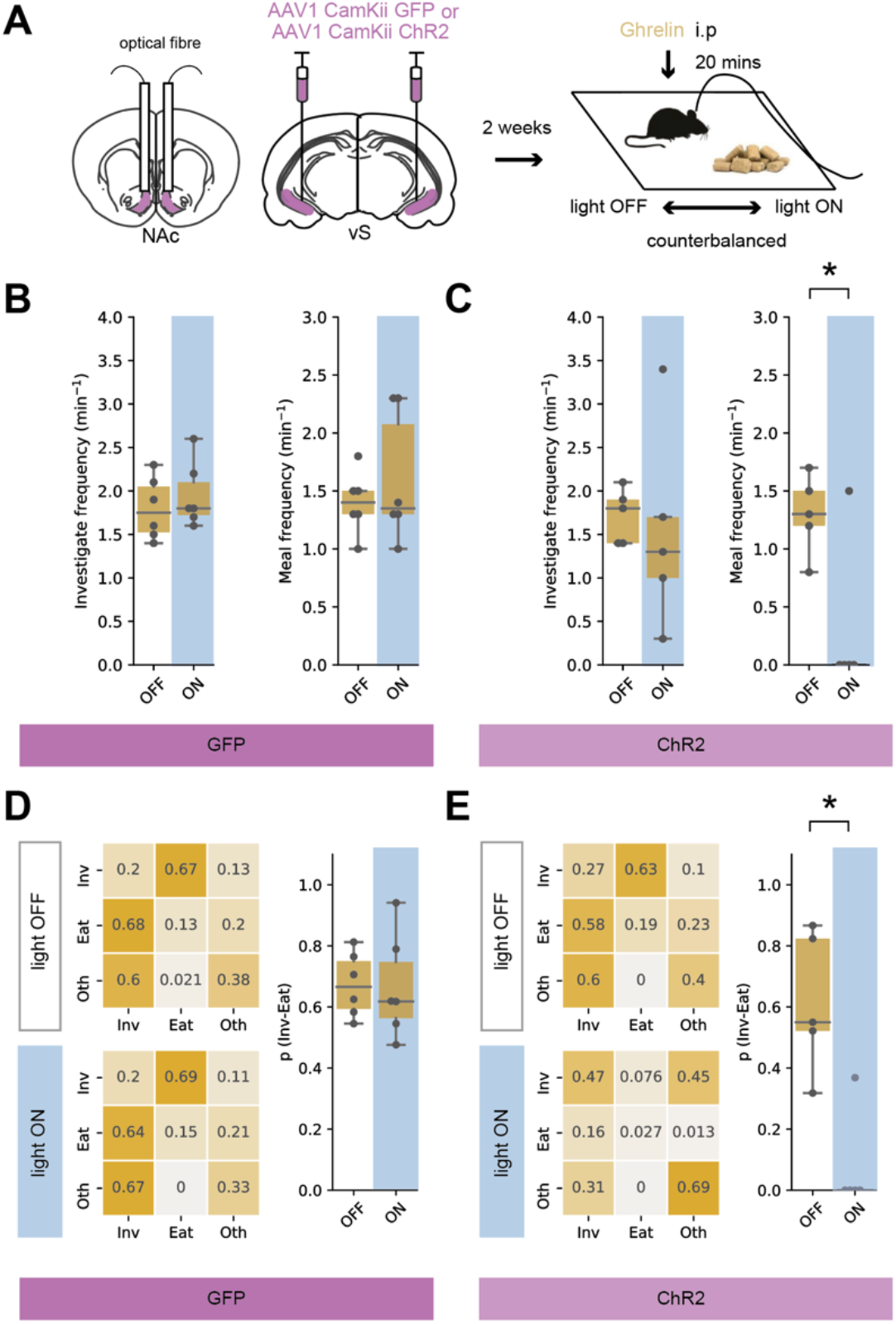
Optogenetic activation of vS-NAc neurons blocks the transition from investigation to eating. **A**. *Left*, schematic of injections allowing optogenetic activation of vS-NAc neurons. *Right*, schematic of experiment. Activity of vS-NAc neurons was manipulated while mice explored a familiar arena containing chow after i.p. injection of ghrelin. For both GFP and ChR2 groups mice underwent the session twice, with 20 Hz light stimulation or no light stimulation in a counterbalanced order. **B**,**C**. Analysis of frequency of investigation and eating in GFP (**B**), and ChR2 (**C**) mice with or without light stimulation. Note that light has no effect on GFP expressing animals. In ChR2 expressing animals there is no effect on investigation frequency, but stimulation markedly reduces frequency of eating. **D**,**E**. Markov analysis of feeding behaviour during ten-minute session in GFP (**D**), and ChR2 (**E**) mice with or without light stimulation. *Left*, state transitions for light off and light on sessions. *Right*, summary of investigation to eat transition across all mice with and without light. Note light has no effect in GFP expressing mice, but results in a large decrease in p(inv->eat) in ChR2 expressing mice.

### Artificially increasing vS-NAc activity blocks ghrelin mediated increases in feeding

Our results so far suggest a model where a high level of eating after increases in peripheral ghrelin is associated with inhibition of vS-NAc activity during food investigation. This is reminiscent of a much-hypothesised role for vS-NAc neurons in behavioural inhibition (Gray and McNaughton, 2003) – where increased activity of vS-NAc neurons inhibits ongoing behaviour (O’Connor et al., 2015; Reed et al., 2018).

We reasoned that if activity of vS-NAc neurons specifically inhibited the transition from investigation to eating, artificial activation of vS-NAc neurons should block ghrelin-induced increases in feeding, but have little effect on other behaviours, in particular the frequency of investigation of food. To test this, we used injections of AAV to express the excitatory opsin channelrhodopsin2 (ChR2) bilaterally in excitatory neurons in vS, and bilaterally implanted optical fibres in NAc (**Figure 4A**). This allowed us to activate vS terminals in NAc with brief pulses of blue light. We then compared mice expressing ChR2 with control mice expressing GFP. We repeated the ten-minute feeding assay in a counterbalanced design where in both cases the mouse was given ghrelin administration, but on one day the mouse underwent 20 Hz blue light stimulation during the session, while on the alternate day no light was present (**Figure 4A**).

In control animals, there was no effect of blue light stimulation, and ghrelin resulted in robust feeding behaviour in both light ON and light OFF days (as seen by high chow consumption, high frequency of investigation of food, and high frequency of eating, **Figure 4B**). Similarly, by creating transition matrices for each animal in each condition, we found that there was a high probability of transitioning from investigation to eating p(Inv->eat), and this was unaltered by light stimulation (**Figure 4D**). However, light delivery in ChR2-expressing animals caused large but specific changes in behaviour. While light stimulation had no influence on the frequency of food investigation, it resulted in an almost complete cessation of eating (**Figure 4C**). Again, by constructing transition matrices for these animals we found that this was due to a marked reduction in p(Inv->eat) (**Figure 4E**). Together, these results suggest that activation of vS-NAc neurons blocks the transition from food investigation to eating, even in the presence of high levels of peripheral ghrelin. This suggests that a key effect of peripheral ghrelin in vS may be to inhibit the activity of vS-NAc neurons, to overcome a ‘block’ this activity imposes on feeding behaviour.

### Increasing peripheral ghrelin increases inhibitory postsynaptic amplitude in vS-NAc neurons

Our results so far suggest that ghrelin may influence hippocampal circuitry by inhibiting the activity of vS-NAc neurons during food investigation. We next wanted to look for the cellular underpinnings of this change. We hypothesised that this decrease in activity may be due to plasticity of inhibitory connectivity. To test this, we injected mice with two colours of retrobeads – one in NAc and one in LH (AlSubaie et al., 2021; Sánchez-Bellot et al., 2022; Wee and MacAskill, 2020). This allowed us, two weeks later, to inject either PBS or ghrelin i.p and after 20 minutes prepare acute slices where we could visualise and record from NAc and LH-projecting vS neurons in acute slices from the same mice (**Figure 5A**).

**Figure 5.**
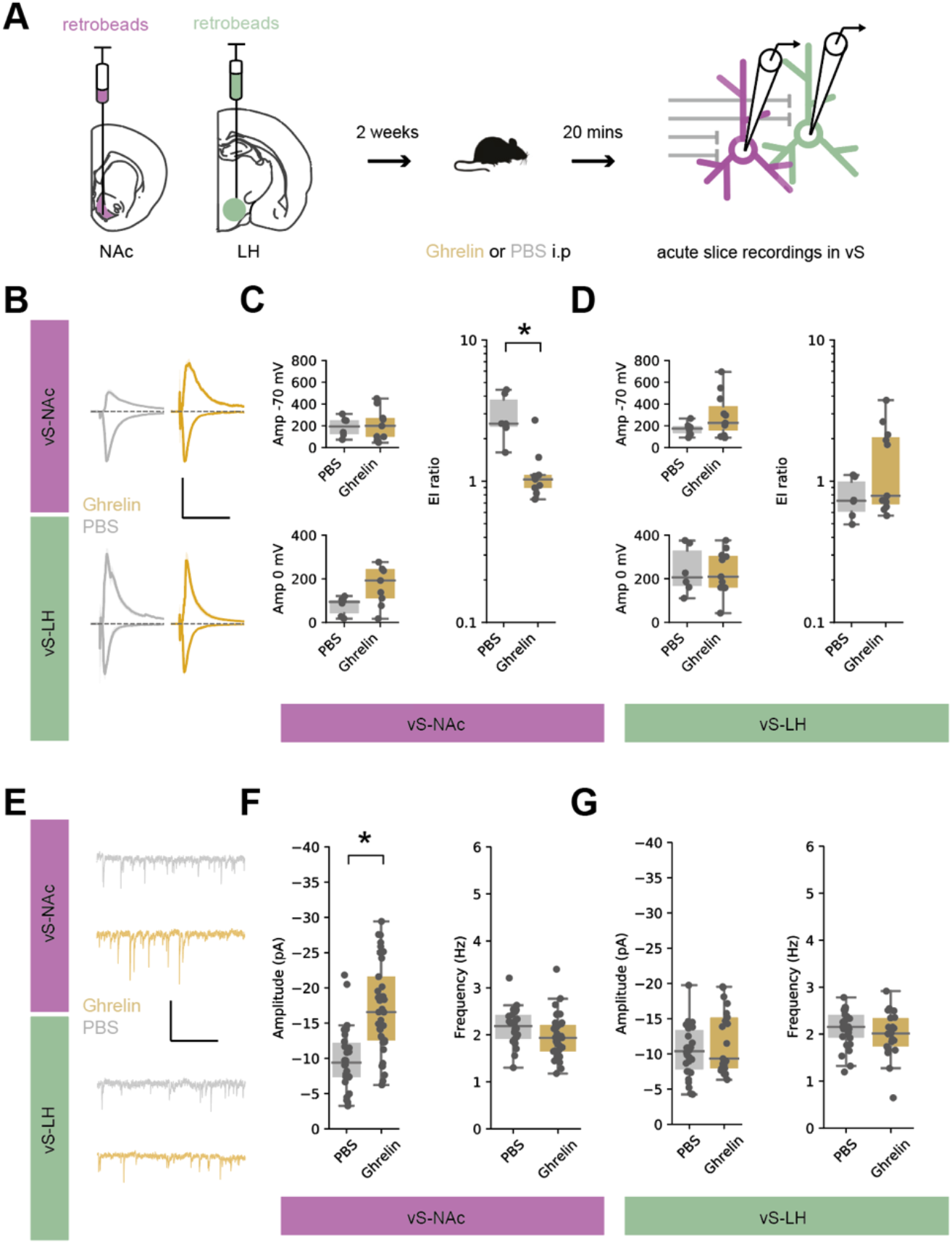
Ghrelin increases the amplitude of postsynaptic inhibition in vS-NAc neurons. **A.**Schematic of retrobead injections allowing whole cell recordings of vS-NAc and vS-LH neurons. Two weeks after injections mice were treated with either ghrelin or PBS control, and acute slices were prepared 20 mins later. **B**. Average electrically evoked PSCs at -70 mV (downwards trace – a proxy for excitatory PSCs) and 0 mV (upwards trace – a proxy for inhibitory PSCs) in vS-NAc and vS-LH neurons after PBS (grey) or ghrelin (gold) treatment. Note increase in amplitude of vS-NAc IPSC after ghrelin treatment. Scale bar = 200 pA, 100 ms. **C**. Left, summary of PSC amplitude in vS-NAc at -70 mV (top) and 0 mV (bottom). Right, summary of E:I ratio (amplitude at -70 mV / amplitude at 0 mV – a proxy for relative excitatory vs inhibitory drive). Note consistent decrease in E:I ratio after ghrelin, suggesting an increase in relative inhibitory drive. **D**. As in (**C**) but for vS-LH neurons. Note lack of consistent changes across any measures. **E**. Example traces containing isolated mIPSCs in vS-NAc and vS-LH neurons after PBS (grey) or ghrelin (gold). Scale bar = 20 pA, 1s. **F**. Summary of amplitude (left) and frequency (right) of mIPSCs in vS-NAc neurons. Note consistent increase in mIPSC amplitude after ghrelin, with no change in frequency. **G**. As in (**F**) but for vS-LH neurons. Note lack of consistent changes across any measures.

We first compared the relative excitatory and inhibitory synaptic strength onto each projection population by calculating the E:I ratio in response to electrical stimulation of the Schaffer collateral input (the ratio of the excitatory current at -70 mV divided by the feedforward inhibitory current at 0 mV, **Figure 5B**). Interestingly we found that while ghrelin administration had no influence on E:I ratio in vS-LH neurons, in vS-NAc neurons there was a large decrease in E:I ratio in mice treated with ghrelin compared to controls, suggesting an increase in relative inhibitory synaptic strength (**Figure 5C, D**).

Next, we recorded miniature inhibitory post synaptic currents (mIPSCs) from both vS-NAc and vS-LH neurons in control and ghrelin treated mice (**Figure 5E**). In these recordings the amplitude of mIPSCs is proportional to the postsynaptic efficacy, while the frequency of events is proportional to both the probability of release and the number of synaptic connections. Consistent with our results above, we found that ghrelin resulted in a large increase in inhibition in vS-NAc neurons, but no changes in vS-LH neurons (**Figure 5F,G**). This increased inhibition was due to an increase in the amplitude of mIPSCs in vS-NAc neurons, with no change in their frequency (**Figure 5F**). Together, this suggests that increased peripheral ghrelin increases synaptic inhibition onto vS-NAc neurons, through an increase in the postsynaptic strength of inhibitory synapses.

### GHSR1a expression in vS-NAc neurons is required for ghrelin mediated changes in inhibitory synaptic strength

We next wanted to understand the mechanism by which peripheral ghrelin can influence synaptic inhibition in vS-NAc neurons. vS neurons express the ghrelin receptor GHSR1a, and peripheral ghrelin is known to cross the blood-brain-barrier and enter the hippocampus, where it has the ability to directly influence postsynaptic properties to influence behaviour (Diano et al., 2006; Hsu et al., 2015, 2016; Kanoski et al., 2013; Ribeiro et al., 2014). Therefore, we wanted to ask if the influence of peripheral ghrelin we observed on vS inhibitory synaptic properties might be via this direct activation of the GHSR1a receptor on the postsynaptic membrane of vS-NAc neurons.

To test this, we developed an AAV containing a cre-dependent RNA interference (RNAi) cassette that robustly knocked down expression of GHSR1a (**Supp Fig 5**). We then used an intersectional viral method as before to reduce levels of GHSR1a only in vS-NAc neurons (**Figure 6A, Supp Fig 5**), and used an RNAi targeted to a scrambled sequence not present in the genome as a control. Thus in these animals, GHSR1a is knocked down only in vS neurons that project to NAc, allowing us to investigate the influence of GHSR1a in these neurons, while leaving GHSR1a expression in other canonical regions such as the arcuate nucleus and LH intact. After allowing 2 weeks for expression, we then prepared acute slices from these animals 20 minutes after i.p. injection with either PBS or ghrelin as before. We then recorded mIPSCs from identified vS-NAc neurons expressing the GHSR1a or control RNAi constructs.

**Figure 6.**
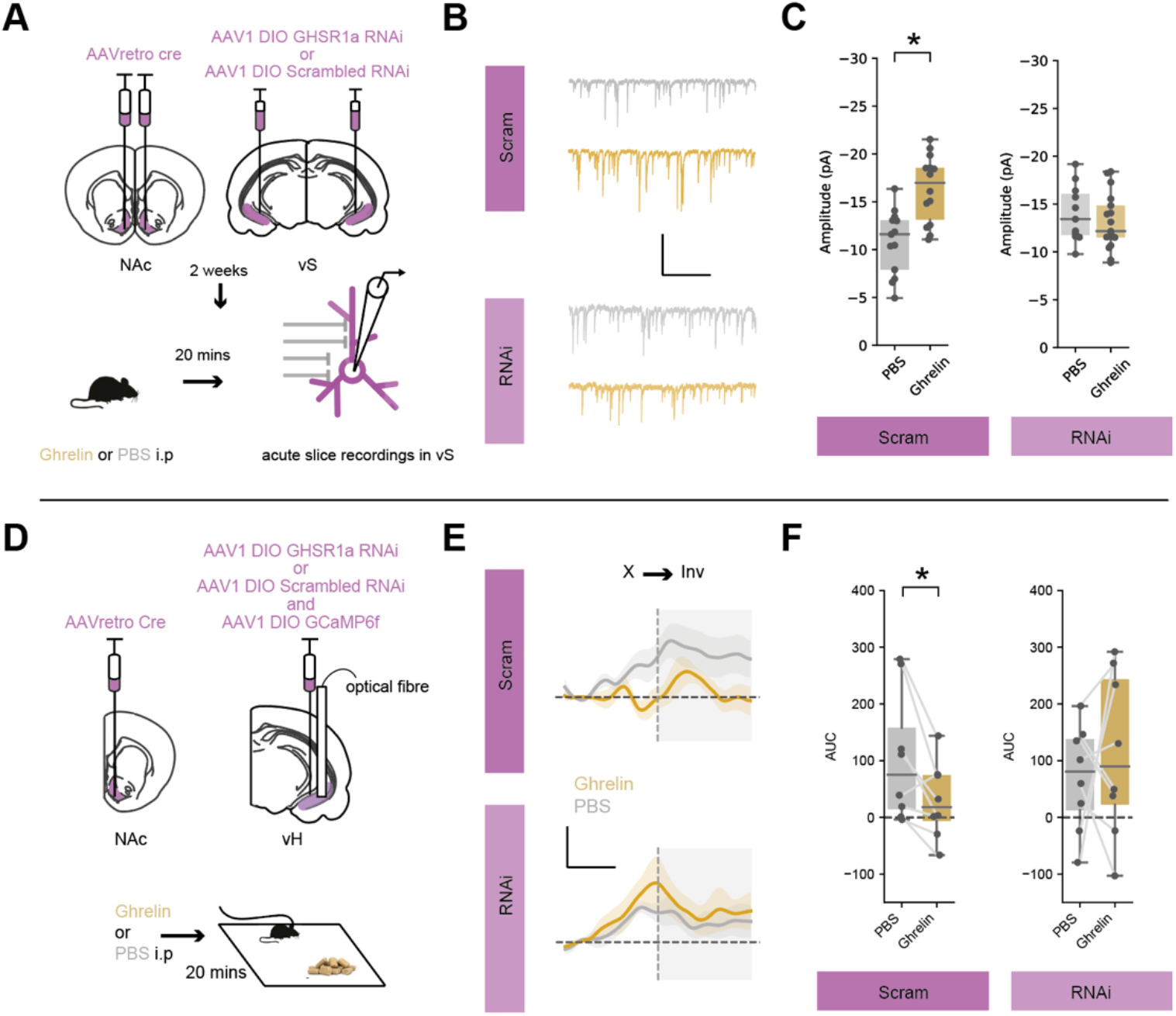
Postsynaptic expression of the ghrelin receptor GHSR1a is required for inhibition of vS-NAc neurons. **A**. Top, schematic of injections allowing intersectional targeting of vS-NAc neurons with either GHSR1a RNAi or scrambled control. Bottom, two weeks after injections mice were treated with either ghrelin or PBS control, and acute slices were prepared for fluorescently identified whole cell recordings 20 mins later. **B**. Example traces containing isolated mIPSCs in vS-NAc neurons expressing GHSR1a RNAi or scrambled control, after PBS (grey) or ghrelin (gold). Scale bar = 15 pA, 1 s. **C**. Summary of amplitude of mIPSCs in vS-NAc neurons. Note consistent increase in mIPSC amplitude after ghrelin in scrambled control neurons, with no change in GHSR1a RNAi neurons. **D**. Top, schematic of injections allowing intersectional targeting of vS-NAc neurons with either GHSR1a RNAi or scrambled control, as well as GCaMP6f and an optical fibre to allow photometry recordings. Bottom, schematic of experiment. Activity of vS-NAc neurons was recorded while mice explored a familiar arena containing chow. **E**. Average activity of vS-NAc neurons across all mice expressing GHSR1a RNAi or scrambled control, aligned to end of food investigation, after injection of either PBS (grey) or ghrelin (gold). Note large increase in activity ramping to end of investigative bout after PBS, but lack of activity after ghrelin in controls, while in GHSR1a expressing mice, activity is high in both PBS and ghrelin. Scale bar = 0.5 zF, 4 s. **F**. Summary of activity around investigation for vS-NAc neurons expressing GHSR1a RNAi or scrambled control, after PBS (grey) or ghrelin (gold). Note increase in activity around investigation is reduced after ghrelin in control mice, while there is no effect of ghrelin in GHSR1a RNAi mice.

Consistent with our previous results, we found that in mice expressing the scrambled control RNAi in vS-NAc neurons, ghrelin administration resulted in an increase in mIPSC amplitude (**Figure 6B, C**). However, in neurons with knockdown of GHSR1a, mIPSCs were almost completely insensitive to ghrelin administration (**Figure 6B,C**). Therefore the changes in inhibitory synaptic connectivity in vS-NAc neurons on administration of peripheral ghrelin require expression of the GHSR1a receptor on the postsynaptic membrane of vS-NAc neurons.

### vS-NAc GHSR1a expression is required for peripheral ghrelin induced alterations in vS-NAc activity during feeding

Our results above suggested that increased synaptic inhibition in vS-NAc neurons after ghrelin treatment is mediated via postsynaptic GHSR1a. Our model proposes that the reduction of vS-NAc activity during investigation of food induced by ghrelin is due to this increased inhibitory drive. If this were true, then GHSR1a knockdown should also block this ghrelin mediated reduction in activity. To test this, we again used intersectional viral infection to unilaterally express either control or GHSR1a RNAi constructs in vS-NAc neurons (**Figure 6D**). In each mouse we also expressed GCaMP6f in vS-NAc neurons, and implanted an optical fibre just above vS. Importantly, due to redundancy across hemispheres, unilateral expression of the RNAi constructs ensured that behaviour was not affected by the manipulation of GHSR1a levels (**Supp Fig 5**), and so allowed investigation of alterations in vS-NAc activity under equivalent behavioural conditions. Together, this approach allowed us to use fibre photometry to monitor the activity of vS-NAc neurons with or without GHSR1a knockdown during feeding behaviour, and to investigate the influence of peripheral ghrelin administration on this activity.

We first found that mice expressing the control RNAi again showed a ramp of activity around investigation in sated mice, and that this was reduced after ghrelin administration (**Figure 6E, F**). In contrast, the activity of vS-NAc neurons in GHSR1a knockdown animals during food investigation remained high even after ghrelin administration (**Figure 6E, F**). Together this suggests that GHSR1a in vS-NAc neurons is required for peripheral ghrelin to increase inhibitory synaptic strength in vS-NAc neurons, and that removal of GHSR1a renders vS-NAc activity during investigation of food insensitive to peripheral ghrelin.

### vS-NAc GHSR1a expression is required for peripheral ghrelin-mediated increase in the transition from investigation to eating

Our results suggest that GHSR1a is required for peripheral ghrelin to inhibit the activity of vS-NAc neurons around food investigation. Moreover, our optogenetic experiments (**Figure 4**) show that this reduction in vS-NAc activity is necessary for ghrelin-mediated increases in transitioning from food investigation to food consumption. Therefore, we hypothesised that GHSR1a in vS-NAc neurons would be a key factor in ghrelin-mediated changes in feeding behaviour. To test this, we intersectionally expressed GHSR1a RNAi bilaterally in vS-NAc neurons, and compared to littermate controls with control RNAi expression (**Figure 7A**). In comparison to our previous unilateral knockdown experiments designed to not affect behaviour, this bilateral expression in this experiment does not allow for compensation by the non-manipulated hemisphere, and can therefore be used to probe behavioural function (in contrast to **Supp Fig 5**). We then used this approach to investigate the influence of GHSR1a knockdown on feeding in response to peripheral ghrelin.

**Figure 7.**
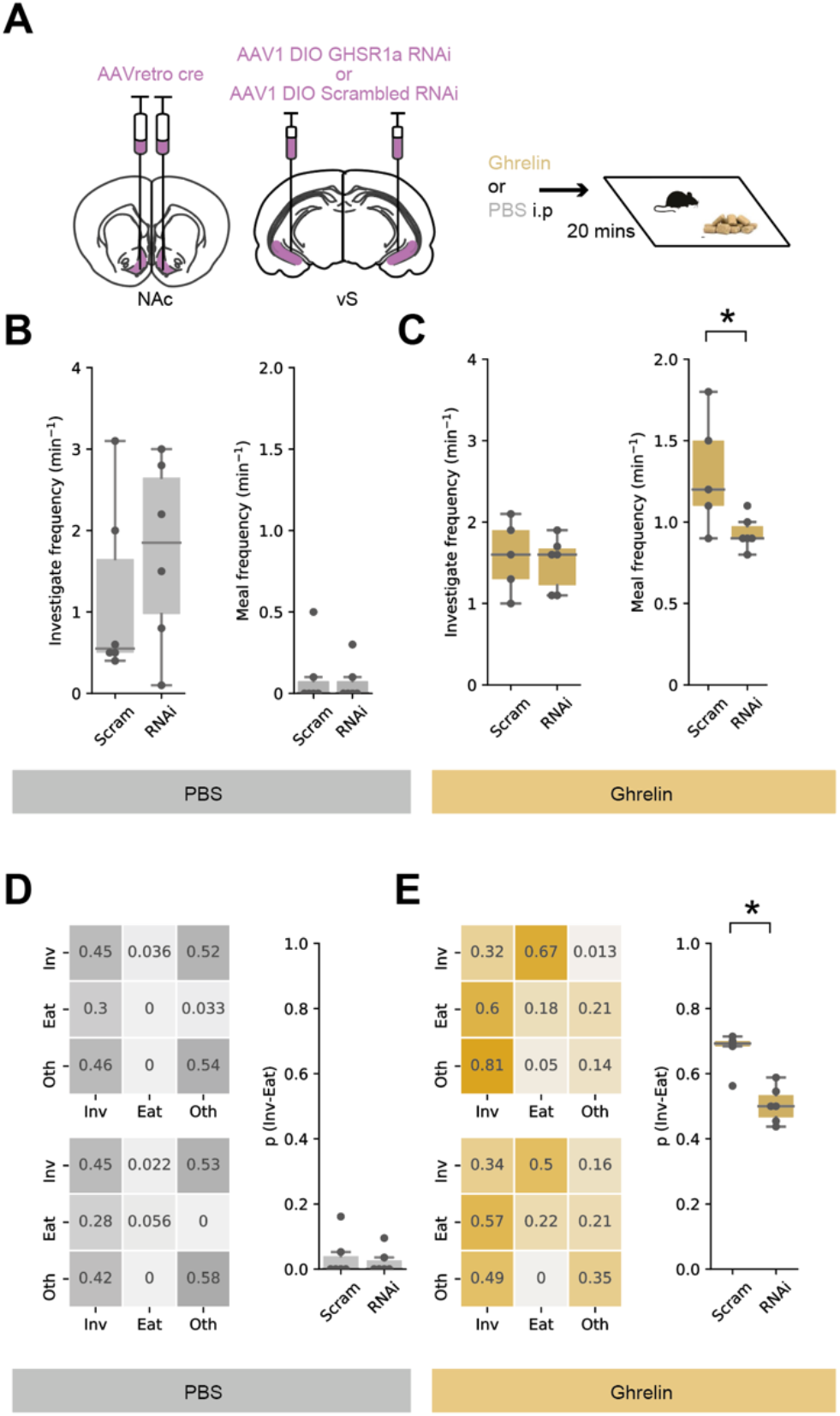
Postsynaptic GHSR1a expression in vS-NAc neurons is required for ghrelin induced increase in the transition from food investigation to eating. **A**. Left, schematic of injections allowing bilateral, intersectional targeting of vS-NAc neurons with either GHSR1a RNAi or scrambled control. Right, schematic of experiment. Mice explored a familiar arena containing chow after an injection of ghrelin or PBS control in a counterbalanced order **B**,**C**. Analysis of frequency of eating, and duration of each eating bout in PBS (**B**), and ghrelin (**C**) treated mice with GHSR1a RNAi or control. Note that RNAi mice are equivalent to control mice in PBS conditions. In contrast after ghrelin, RNAi mice show reduced frequency of eating. This decrease was accompanied by no change in eating duration. **D**,**E**. Markov analysis of feeding behaviour during ten-minute session in PBS (**D**), and ghrelin (**E**) treated mice with GHSR1a RNAi or control. Left, state transitions for Control (top) and RNAi (bottom). Right, summary of investigation to eat transition across all mice. Note RNAi has no effect on PBS treated mice, but results in a decrease in p(inv->eat) in after ghrelin treatment.

We first looked at the effect of GHSR1a knockdown on the frequency of feeding associated behaviours. We found no effect of GHSR1a knockdown on either investigation of food, or feeding in PBS treated mice (**Figure 7B**). This suggests that there is little role for constitutive GHSR1a activity in sated mice, potentially due to a strong floor effect (Kern et al., 2015). After ghrelin treatment, control animals maintained similar levels of food investigation, but this was accompanied by a large increase in the frequency of eating (**Figure 7C**). In contrast, while GHSR1a RNAi animals showed a similar level of investigation of food, this was accompanied by a reduction in the frequency of eating compared to control animals – similar to the effect of artificially activating vS-NAc neurons with ChR2 (**Figure 4**). Next, we constructed transition matrices for each animal in each condition, and again saw a consistent and specific reduction of the probability to transition from investigation to eating p(Inv->eat) (**Figure 7D, E**).

Overall, we have shown that peripheral ghrelin increases inhibitory synaptic strength onto vS-NAc neurons via a mechanism that requires GHSR1a. This increased inhibition accompanies reduced vS-NAc activity during investigation of food, which promotes the transition from investigation to eating. This mechanism is consistent with a classically described role for the ventral hippocampus in behavioural inhibition (Gray and McNaughton, 2003), and provides a cellular basis of hippocampal resolution of the conditional ambiguity of food cues (Davidson et al., 2007; Gershman, 2017). Moreover, this provides the circuit basis for the direct interaction of cognitive and goal directed brain regions in the control of feeding, and provides a mechanistic locus for the close interaction between diet, health and cognitive ability.

## DISCUSSION

In this study, we have shown a key circuit mechanism for how peripheral ghrelin dramatically alters feeding behaviour. We observed large increases in vS-NAc activity during investigation of food, and this activity inhibited the transition from food investigation to consumption. Consistent with a key role, we found that increases in peripheral ghrelin act via modulation of this activity through postsynaptic GHSR1a signalling. Via this interaction, peripheral ghrelin increases the amplitude of postsynaptic inhibition, reducing vS-NAc activity during food investigation, and thus facilitating an increase in the transition to eating. The identification of a ghrelin-sensitive vS circuit that mediates food-seeking behaviour provides one of the first mechanistic descriptions of how the hippocampus may sense an internal state like hunger *in vivo*.

### The role of ghrelin in anticipatory versus consummatory behaviour

Our data demonstrate that ghrelin inhibits vS activity specifically during food anticipation, and suggest that it is this inhibition that promotes the transition to eating. On the surface, this goes against classic notions of ghrelin as a ’hunger hormone’ that mimics fasting by increasing food consumption (Tschöp et al., 2000; Wren et al., 2001). However, this is consistent with several observations suggesting ghrelin mediates food anticipation rather than food consumption. For example, the concentration of circulating ghrelin accurately reflects the anticipation of an upcoming meal (Cummings et al., 2004; Drazen et al., 2006), and studies employing pharmacological and genetic disruption of GHSR1a, both systemically and in vS, show that mice have disrupted anticipatory behaviour preceding a meal (Blum et al., 2009; Kanoski et al., 2013; Verhagen et al., 2011). Furthermore, GHSR1a-null mice have normal body weight (Sun et al., 2008), indicating that ghrelin is dispensable in homeostatic food intake regulation. Importantly, we did observe increases in the duration of each eating bout in ghrelin treated animals, consistent with classic descriptions of ghrelin function (**Sup Figure 1**). However, only the frequency of meal initiation was altered by vS-NAc GHSR1a manipulations (**Figure 7C**), with meal duration unaffected (**Sup Figure 6**). This dissociation of function suggests that multiple roles of ghrelin may be separated across distinct regions and circuits – while the role of ghrelin in vS seems to be specifically in anticipation of feeding, different effects of ghrelin may be mediated by other brain regions such as hindbrain or hypothalamus (Rossi and Stuber, 2018).

### Mechanisms of ghrelin signalling in vS

Given the tight regulation of substance entry across the blood-brain barrier (BBB), one important question is whether ghrelin and other hormones mediate their effects on vS circuitry by directly binding hippocampal neurons, or are instead signalled indirectly via upstream synaptic inputs that themselves have access to peripheral ghrelin. The hippocampus is situated adjacent to circulating cerebrospinal fluid (CSF) in the ventricles, and has a rich surrounding blood supply from the choroid plexus (Lathe, 2001) that facilitates transfer of molecules across the BBB (Hamasaki et al., 2020), including ghrelin (Banks et al., 2002; Diano et al., 2006). This is consistent with a vast array of peripheral ghrelin mediated structural and functional effects on hippocampal neurons (Diano et al., 2006; Ribeiro et al., 2014), but also the role of hippocampal ghrelin in influencing behaviour (Diano et al., 2006; Hsu et al., 2015; Kanoski et al., 2013). More generally, the hippocampus expresses functional receptors for a multitude of peptide hormones similar to ghrelin (Lathe, 2001), such as leptin (Irving and Harvey, 2014) and insulin (Soto et al., 2019). Together this suggests that these hormones are likely capable of binding to hippocampal neurons to affect their function.

It is important to note however that the influence of ghrelin of vS circuitry may occur via other, indirect mechanisms. First there is a line of evidence that proposes that peptide hormones like ghrelin, insulin and leptin are unable to cross the BBB (Furness et al., 2011; Kern et al., 2015) without specialised mechanisms (Banks et al., 2002), and it is unclear if these are present in vS. While the contradictory findings surrounding ghrelin access to vS could be explained by BBB permeability being extremely plastic (for example, accessibility of ghrelin is itself often dependent on hunger state (Banks et al., 2008; Langlet et al., 2013)) there remains a possibility that direct permeability through the BBB might not be the major route for ghrelin to influence vS.

As a result, multiple alternate mechanisms have been proposed to explain the role of GHSR1a signalling in vS. First, it has been proposed that the ghrelin receptor GHSR1a exerts its effect through ligand independent constitutive activity (Petersen et al., 2009). However, this is hard to reconcile with past experiments showing the presence of peripheral ghrelin in hippocampus (Diano et al., 2006), as well as other experiments (Hsu et al., 2015; Kanoski et al., 2013) including ours (**Figure 5, 6** and **7**) showing an active role of the receptor in response to stimuli, suggesting a role beyond constitutive activity. There may therefore, be a combination of ligand-dependent and ligand-independent activity within the GHSR1a system in vS. A key point of future work will be to understand the distinct roles of these two potential mechanisms.

A second alternative to direct ghrelin sensing in vS, is through dimerization of the GHSR1a receptor with the D1 dopamine receptor (Kern et al., 2015), which could render vS dopamine signalling dependent upon GHSR1a expression. In this scenario, ghrelin could indirectly activate ghrelin-sensing LH neurons that then promote VTA activity (Cone et al., 2014, 2016). This activity may then provide a dopaminergic input to hippocampus that through dimerisation may be sensitive to the presence GHSR1a. This also has the potential to be the case for other neuromodulators or neuropeptides that could also act as co-agonists to ghrelin signalling in vS, including acetyl-choline from septal areas, serotonin from raphe or melanin concentrating hormone from hypothalamus (Colgin et al., 2003; Noble et al., 2019; Yang et al., 2020).

Together, while our study shows that vS-NAc circuitry is sensitive to the level of peripheral ghrelin, and this sensitivity is dependent on postsynaptic expression of the ghrelin receptor GHSR1a, future work is needed to explore whether ghrelin directly modulates hippocampal activity via BBB permeability or indirectly through modulation of upstream synaptic inputs.

### The role of vS in hunger-sensitive goal directed behaviour

It is becoming increasingly understood that the hippocampus not only encodes the relationships between distinct cues in the environment (Eichenbaum, 2017; Rudy and Sutherland, 1995), but also the value of outcomes (Knudsen and Wallis, 2021; Lee et al., 2012), and their interaction. For example, the hippocampal-to-NAc projection has been proposed to be important for the learning of value associations in both physical and abstract dimensions (Duncan et al., 2018; Ito et al., 2008; LeGates et al., 2018; Trouche et al., 2019), and to relay these signals to ventral striatum (AlSubaie et al., 2021; MacAskill et al., 2012; Pennartz et al., 2011; Trouche et al., 2019). This ability is proposed to allow the utilisation of past experience to anticipate the outcome of upcoming behaviour. Consistent with this proposed role, we found that vS-NAc was specifically active around food investigation, and that this activity ramps up during investigative behaviour, consistent with studies of spatial goal directed navigation (Ciocchi et al., 2015). This activity was anticorrelated with the value of the anticipated outcome (Ciocchi et al., 2015; Lafferty et al., 2020; Reed et al., 2018) and the overall amount of food consumed during the session (Yang et al., 2020) – when sated, vS-NAc activity was high, and this correlated with a low probability of initiating eating (**Figure 3**). In contrast, when peripheral ghrelin was high, vS-NAc activity was low. Interestingly, this reduction in anticipatory vS-NAc activity during investigation was distinct from a separate decrease in activity upon food consumption that was insensitive to ghrelin (**Figure 3**, Reed et al., 2018). This is consistent with vS ghrelin sensing only influencing the initiation of eating, but not the increase in eating duration that also results from increases in peripheral ghrelin (**Supp Fig 6**). Therefore, while vS-NAc activity is responsive to the initiation of eating (Reed et al., 2018), and manipulation of vS activity can directly influence consummatory behaviour (Reed et al., 2018, Yang et al., 2020), ghrelin signalling in vS influences only the anticipatory activity leading up to food consumption.

Downstream of vS, our work aligns well with a proposed circuit mechanism for the rapid control of feeding behaviour promoted by activity of D1 dopamine receptor expressing medium spiny neurons (D1 MSNs) in NAc (Lafferty et al., 2020; O’Connor et al., 2015; Reed et al., 2018). vS input to NAc preferentially targets D1 MSNs (MacAskill et al., 2012, 2014), activity of which can rapidly supress reward consumption, and promote exploratory and seeking behaviours (Britt et al., 2012, Reed et al., 2018, O’Connor et al., 2015). In particular, during consummatory behaviour, a subpopulation of D1 MSNs that project to LH directly inhibit eating via targeting of LH GABAergic neurons (Hsu et al., 2015; O’Connor et al., 2015). Thus, through this specialised connectivity, vS-NAc ghrelin sensitivity is ideally situated to provide strong and acute control over feeding behaviour, and provides a means for this control to be utilised to shape behaviour based on internal state.

### Internal state as a distinct dimension of the hippocampal representation

The hippocampus has long been viewed as circuit by which otherwise ambiguous cues can be separated through their association with other cues that occur in close spatial and temporal proximity (Per Andersen et al., 2006). While such representations are often studied in terms of sensory perceptions such as vision audition and olfaction, our work complements a number of studies suggesting that internal state may also be used to disambiguate such situations (Azevedo et al., 2019; Carey et al., 2019; Davidson and Jarrard, 1993; Davidson et al., 2007; Kennedy and Shapiro, 2009; Min et al., 2011; Mohammad et al., 2021; Wallner-Liebmann et al., 2010; Wang et al., 2006). In this study, we show that ghrelin sensing in vS has a key role in modulating behaviour towards food dependent on internal state. However, it is likely that ghrelin sensing in the hippocampus is integrated with other modalities in order to be utilised more generally - for example, to allow the balance of approach and avoidance behaviour, and passive and active strategies in potentially threatening situations (Davidson et al., 2010; Sánchez-Bellot et al., 2022), and context dependent associative learning more generally (Azevedo et al., 2019; Davidson and Jarrard, 1993; Livneh et al., 2017; Mohammad et al., 2021).

### Conclusion

In summary, we describe a ghrelin-sensitive hippocampal circuit that shapes feeding decisions. This circuit provides a locus for understanding how internal states and sensory stimuli are integrated to support anticipatory goal-directed behaviour and how disruption of this process may give rise to disease (Toates, 1981). More generally, our data are in line with a role of the hippocampus in using multimodal contextual cues to anticipate the outcome of upcoming behavioural choices, and to utilise this information to inform decision making based on past experience (Shadlen and Shohamy, 2016).

## METHODS

### Animals

Young adult C57BL/6J male mice (behavioural and anatomical experiments: at least 7 weeks old; whole-cell electrophysiology experiments: 7 – 9 weeks old) provided by Charles River were used for all experiments. All animals were housed in cages of 2 to 4 in a temperature- and humidity-controlled environment with a 12 h light-dark cycle (lights on at 7 am to 7 pm). Food and water were provided ad libitum. All experiments were approved by the UK Home Office as defined by the Animals (Scientific Procedures) Act, and strictly followed University College London ethical guidelines.

### Viruses

Titre
|  |  |
| --- | --- |
| rAAV2-retro-Syn-Cre | $5.3 \times 10^{12}$ |
| AAV1-CAG-Flex-GCaMP6f-WPRE-SV40 | $>1 \times 10^{13}$ |
| AAV1-CamKII-GFP | $>1 \times 10^{13}$ |
| pAAV-CamKII-hChR2(H134R)-EYFP | $2.5 \times 10^{13}$ |
| AAV8-hSyn-DIO-hM3Dq | $7.9 \times 10^{12}$ |
| AAV8-hSyn-DIO-hM4Di | $7.0 \times 10^{12}$ |
| AAV8-hSyn-DIO-mCherry | $3.8 \times 10^{12}$ |
| AAV1-EF1a-DIO-mCherry-scrmb-shRNAmir | $1.0 \times 10^{13}$ |
| AAV1-EF1a-DIO-mCherry-ghsr-shRNAmir | $4.1 \times 10^{13}$ |

### Stereotaxic surgery

Stereotaxic surgeries were performed according to previously described protocols (Cetin et al., 2006). Mice were anaesthetised with isoflurane (4% induction, 1.5 to 2% maintenance) and secured onto a stereotaxic apparatus (Kopf). A single incision was made along the midline to reveal the skull. AP, ML and DV were measured relative to bregma, and craniotomies were drilled over the injection sites.

Stereotaxic coordinates:

| Region | ML | AP | DV |
| --- | --- | --- | --- |
| Lateral hypothalamus | 0.9 | -1.3 | -5.2 |
| Nucleus accumbens (medial shell) | 0.9 | +1.1 | -4.6 |
| Ventral subiculum | 3.4 | -3.2 | -4.3 |

Long-shaft borosilicate pipettes were pulled and backfilled with mineral oil, and viruses were loaded into the pipettes. Viruses were injected with a Nanoject II (Drummond Scientific) at a rate of 13.8 to 27.6 nL every 10 s. Following infusion of the virus, the pipette was left in place for an additional 10 mins before being slowly retracted. For anatomy experiments, following injection of substances into the brain, animals were sutured and recovered for 30 mins on a heat pad. Animals received carprofen as a peri-operative s.c. injection (0.5 mg/kg) and in their drinking water (0.05 mg/mL) for 48 hours post-surgery.

For photometry and optogenetic experiments, fibre cannulae were implanted following virus injection in the same surgery. The skull was roughened and two metal screws were inserted into the skull to aid cement attachment. Photometry cannulas were targeted to ventral CA1/subiculum, optogenetic cannulas were inserted at a 10-degree angle targeted to NAc shell. Cannulas were secured to the skull by applying two layers of adhesive dental cement (Superbond CB). The skin was attached to the cured dental cement with Vetbond. Animals received a subcutaneous injection of carprofen (∼5 μL of 0.5 mg/mL stock) prior to recovery in a warm chamber for 1 hour and continued receiving carprofen in their drinking water (0.05 mg/mL) for 48 hours post-surgery. Mice were allowed to recover for a minimum of 3 weeks before starting photometry experiments. For projection-specific expression of GCaMP6f, 150 – 200 nL of rAAV2-retro-Syn-Cre (Tervo et al., 2016) was injected into the output site (LH or NAc); in the same surgery, 300 – 400 nL of a 1:3 dilution of AAV1-CAG-Flex-GCaMP6f-WPRE-SV40 in sterile saline was injected into vS. This dilution protocol was used to delay excessive GCaMP expression, which could lead to reduced Ca2+ variance in the signal, affect cellular processes and reduce cell health (Resendez et al., 2016). For optogenetic experiments, 400 nL of either AAV1-CamKII-hChR2(H134R)-EYFP, or AAV1-CamKII-GFP as a control were injected into vS. For combined projection-specific fibre photometry and molecular knockdown experiments, 150 – 200 nL rAAV2-retro-Syn-Cre was injected into NAc, and a 1:1 mix (400 nL) of AAV1-CAG-Flex-GCaMP6f-WPRE-SV40 and AAV1-EF1a-DIO-mCherry-ghsr-shRNAmir or AAV1-EF1a-DIO-mCherry-scrmb-shRNAmir was injected into vS. For bilateral knockdown of GHSR1a, 200 nL rAAV2-retro-Syn-Cre was injected into NAc, and 800 nL of AAV1-EF1a-DIO-mCherry-ghsr-shRNAmir or AAV1-EF1a-DIO-mCherry-scrmb-shRNAmir were injected into vS.

### Behaviour

Following at least 3 weeks post-surgical recovery, animals (10 – 12 weeks old) were manually handled for at least 7 days before testing. During the last 3 days of habituation, empty plastic weighing boats were provided in the home cage to habituate the animals to these objects. During this time animals were also habituated to intraperitoneal (i.p.) injection, patch cord attachment (for photometry and optogenetic experiments) and the behavioural boxes as described below. All behavioural experiments were carried out in MEDPC sound-attenuating chambers containing behavioural boxes (21.59 × 18.08 × 12.7 cm) with blank walls. Video recordings were conducted with infrared cameras positioned above each chamber, and video was acquired at 15 or 25 Hz using Bonsai (Lopes and Monteiro, 2021). The different frame rates were due to a change of PS4 cameras over the course of experiments, and this difference in frame rates did not affect the resolution of capturing naturalistic behaviour given the relatively slower time course of evolving behaviours during feeding. All experiments were performed consistently during the middle-to-end of the light cycle (from 2 pm to 7 pm) to control for circadian rhythm variables.

For all behavioural experiments, after acclimatisation to handling and the behavioural chambers, mice were habituated over the course of 2 to 3 days with three i.p. injections of 100 μl sterile phosphate-buffered saline (PBS) to habituate them to manual scruffing and i.p. injection. Following this, animals for photometry and optogenetics experiments were habituated to patch cord attachment for a 10-minute period, as described above. At the end of this habituation period, animals were given an i.p. injection of either ghrelin (2 mg/kg; Tocris) or vehicle control (phosphate-buffered saline, PBS; pH = 7.2). The order of the injections was counterbalanced across animals. The volume of the i.p. injection was fixed at 100 μl. Animals were allowed 15 mins to recover post-injection before the presentation of non-food and food objects. The day of ghrelin injection was selected randomly for each animal, and PBS and ghrelin injection days were spaced apart for a duration of at least 24 hours. After termination of each testing session, the amount of chow consumed during the 10 min presentation was weighed; any spillage of food was recovered and subsequently weighed.

For photometry experiments, the time of food or non-food presentation was noted down and used to manually synchronise the photometry recordings to the start of stimulus presentation. Photometry experiments with apparent failure in equipment or software acquisition, photometry signals from experiments in which the signal did not exceed >2 standard deviations above a 50-s preceding baseline before food presentation, or mice with misplacement of or damage caused by injections and implantations were excluded (4 in total).

For optogenetics experiments, light power at the end of each bilateral patch cord was 8 mW. Light delivery was given in a counterbalanced design, separated by at least 48 hours. On stimulation day, light was delivered at 20 Hz (5 ms on, 45 ms off) throughout the duration of the experiment. One mouse was excluded due to mild seizure activity upon light stimulation.

### Annotation of feeding behaviour

Feeding behaviour was analysed as a composite sequence of five simple, distinct and reproducible behaviours. These elemental behaviours were: Investigate (sniffing or tactile interaction with the object or food without eating), Eat (biting food or chewing movements close to food), Rear (standing on hindlegs while elevating head, can be supported on the walls i.e. thigmotaxis), Groom (licking/scratching of fur, limbs or tail, usually high-frequency movement) and Rest (motionless, usually in corner of box). These behaviours together are referred to as the Behavioural Satiety Sequence (BSS, Halford et al., 1998). These features were manually scored offline using Ethovision XT13 (Noldus). Where possible, manual scoring was conducted in a blinded fashion to experimental groups. For a subset of videos, two independent scorers conducted manual annotation of the behaviour videos to ensure reproducibility. Manual annotation of BSS behaviours from 10-minute videos spanning the food or non-food object presentation period were conducted at 15 or 25 Hz on a frame-by-frame basis. This manual annotation produced vectors of 0s and 1s, where 0 indicates the absence and 1 the presence of the BSS behaviour.

### Analysis of feeding behaviour as a stochastic Markov process

Each behavioural dataset exists as a sequence of BSS behaviours. In other words, the behaviour for a given animal is described by a vector of BSS behaviours occurring over time. Although the total time spent engaging in one behaviour can be computed from this vector, additional information regarding an animal’s feeding strategy exists in the sequence of expression of each BSS behaviour (Burnett et al., 2019). To analyse this sequential information in more detail, we analysed the annotated behavioural patterns for each mouse as a stochastic Markov process that defined the animal’s feeding strategy when presented with chow across different states of hunger. Specifically, a Markov chain is a vector of states that change as a function of time. In this case, the Markov chain is comprised of 5 states corresponding to the 5 BSS behaviours. These Markov chains are described fully by a transition matrix P, where the P*ij* term represents the transition probability from BSS behaviour *i* to *j*. As there are 5 BSS behaviours, P is a 5 × 5 transition matrix, where the rows represent the current BSS behaviour, the columns represent the BSS behaviour one-step ahead and the values in each row sums to 1. For display purposes, as non feeding behaviours showed little consistent, but independent alterations upon ghrelin injection (**Supp Fig 1**), we constructed simpler, 3 × 3 transition matrices by combining non-feeding behaviours into a single state. For comparison, 5 × 5 matrix analysis is provided in **Supp Fig 2**. To compute the empirical transition matrices for each animal, the frequency of each possible transition from behaviour *i* was calculated and normalised by the total number of behavioural transitions occurring from behaviour *i*. These transitions are assumed to be Markovian, which simplifies the calculation of the transition probability P(state = j | state = i). Specifically, the probability of transitioning from state i to state j is dependent only on the current state i and not on states preceding state i. For each animal, there were two transition matrices, *P*_PBS_ and *P*_Ghrelin_. Importantly, these Markov chain vectors disregarded information relating to duration, i.e. the time spent in engaging in one behaviour and the inter-event duration. In other words, by focusing on transitions between BSS behaviours, this analysis was conducted time-agnostically; this method has been shown to accurately capture moment-to-moment behavioural strategies under differing contexts (Burnett et al., 2019) by focusing on the transition probability from one behavioural bout to the next. For example, the vector [Investigate, Investigate, Eat, Groom] represents four distinct BSS bouts of variable length within and between bouts, but only the transitions between bouts were analysed. Note behavioural data in **Figure 1** contains mice also utilised for photometry experiments in **Figure 2** and **3**.

### Analysis of Ca2+ signals from fibre photometry

Measurement of calcium fluorescence signals was carried out as detailed previously (Lerner et al., 2015; Sánchez-Bellot et al., 2022). 470 nm and 405 nm LEDs were used as excitation sources, and the light amplitudes were modulated sinusoidally at 500 Hz and 210 Hz carrier frequencies, respectively. The excitation light was passed through excitation filters (for 470 nm and for 405 nm wavelengths), and a dichroic mirror to combine the two LEDs into a single beam. A 49/51 beam-splitter was used to split the beam into two independent excitation beams for simultaneous recording of two animals. The excitation light was coupled through a fibre collimation package into a fibre patch cord, and linked to a large core (200 μm), high NA (0.39) implant cannula (Thorlabs). Emitted fluorescence signals were collected through the same fibre. Fluorescence output signal was filtered through a GFP emission filter (transmission above 505 nm) and focused onto a femtowatt photoreceiver. The photoreceiver was sampled at 10 kHz, and each of the two LED signals was independently recovered using quadrature demodulation on a custom-written Labview software: this process involved using an LED channel as a signal reference, taking a 90-degree phase-shifted copy of this reference signal and multiplying these signals in quadrature. The multiplied signal was then low-pass filtered with a Butterworth filter (order = 3, cut-off frequency = 15 Hz). The hypotenuse was then computed using the square root of the sum of squares of the two channels. The result corresponds to the demodulated signal amplitude and was decimated to 500 Hz before storing to disk.

To correct for artefacts resulting from Ca^2+^-independent processes such as movement, the Ca^2+^-independent 405 nm isosbestic wavelength signal was scaled to the 470 nm wavelength. The coefficients for the scaling were computed through a least-squares linear regression between the 405 nm and 470 nm signal. This estimated motion (scaled 405 nm) signal was then subtracted from the 470 nm signal to obtain a pure Ca^2+^-dependent signal.

Calcium activity signals time-locked to the presentation of chow were extracted using the time of presentation manually determined from video recordings. The signal was decimated to 15 Hz, z-score normalised, filtered with a Gaussian filter (σ = 1.5) and baselined to the mean signal in the -50 to -10 seconds preceding the time of presentation of food or non-food object. For event-triggered analysis, the photometry signal was aligned to the onset of each behavioural event obtained from the manually scored behaviour. The behavioural events were clustered into bouts (defined as continuous engagement in the behaviour), and the onset of each bout was taken as the time point to align the photometry signal. A peri-event window of 20 s surrounding the onset of the behaviour was obtained for each signal, and the resulting signal was baselined to the time period from -10 to -7.5 seconds relative to the onset of each event. All trials obtained for an animal were averaged to obtain a nested average event-triggered signal; these signals were then averaged across mice to obtain the population event-triggered average. Due to the stochastic nature of emitting a given behaviour, not all behaviours were present in all animals. To avoid including the same behavioural events across multiple event triggered averages, only events separated by more than 5s were included. Animals displayed Investigative behaviour consistently in all internal states of hunger, but the proportion of animals showing Eat, Groom, Rest and Rear behaviours were variable.

### Linear encoding model relating behaviour to neural activity

To quantify the contribution of each BSS behaviour to neural activity, a multiple regression model was used. The linear model was constructed using the Python package sklearn, with the z-scored baselined photometry signal as the dependent variable, and a regressor matrix of BSS behavioural arrays as independent variables. The regressor matrix contained 27 regressors in total: 5 behavioural regressors (Investigate, Eat, Rear, Groom and Rest), 20 behavioural transition regressors (for example, Investigate → Eat), a manual presentation regressor and a velocity regressor. The 5 behavioural regressors were coded as pulses of 0s and 1s, where 1s indicate the engagement in a BSS behaviour and 0s the absence of engagement. The 20 transition regressors were included to account for any possible contribution of behavioural transitions to the photometry signal, and were derived as follows: a Markov chain vector of BSS behaviours was produced at 15 or 25 Hz and any across-BSS transitions (e.g. Investigate → Eat, not Investigate → Investigate) occurring within 5 seconds was emitted as a temporal pulse of 1 at the onset of the next BSS behaviour. To account for temporal distortion of the behavioural transition period in the associated Ca2+ activity, an exponential function was first computed:

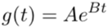

where

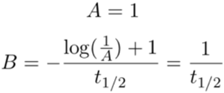

where t1/2 is the half-life of the exponential function and set to 2 seconds. The transition regressor was convolved with the exponential function:

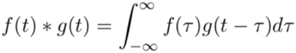

where f(t) is the transition regressor and g(t) is the exponential function. This produces a sharp peak to 1 and a decay rate of t1/2. The exponential decay function was set to have a half-life of 2 s to approximate near-complete decay of the GCaMP6f signal. The presentation regressor was set to an exponential function with a peak time at presentation onset and a decay rate of 5 seconds to capture the salience of stimulus presentation. Finally, the velocity regressor was a continuous variable tracking the instantaneous velocity of the animal derived from position tracking using Ethovision. To handle periods of discontinuous tracking, the velocity data were pre-processed with a ceiling of 5 cm/s (to handle large jumps in tracking), smoothed with a rolling mean filter (window = 3 seconds), and imputed via linear interpolated to handle missing values.

The final linear encoding model was therefore the following:

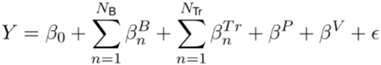

where Y is the dFz in one animal, β0 is the intercept (bias), ε is a Gaussian noise term, NB and NTr are the numbers of the behavioural and transition regressors (5 and 20, respectively), βB, βTr, βP and βV are the beta weights for the behavioural, transition, presentation and velocity regressors, respectively. Specifically, the beta weights βB can be interpreted as the isolated, average neural response to engagement in that BSS behaviour. The crucial aspect of the linear encoding model is the simultaneous inclusion of possible confounding variables, for example, behavioural transitions and velocity, which may each contaminate the neural response. The linear model thus statistically disambiguates the neural response to BSS behaviour engagement from other events in close temporal proximity.

The linear model was fit using ridge regression, a version of the ordinary least-squares regression that penalises the size of the estimated β coefficients by L2 regularisation. This ensures that the β weights were constrained to avoid overfitting, and the penalty term α adjusts the degree of shrinkage of the β weights. Prior to fitting, the dataset was split into an 80% training set to estimate the β weights and 20% test set for evaluating the model predictions. On this training dataset, a nested cross-validation procedure was used: first, the training dataset was split into 5 folds for evaluation. For each fold, the α hyperparameter was tuned using leave-one-out cross-validation (GCV). GCV works analogously to a grid search by exploring the alpha parameter space, and selecting the α value that maximises the prediction accuracy of the model; the values of α tested were 10−3, 10−2, 10−1, 100 and 101, using the function RidgeCV on Python’s sklearn package. The values of α did not differ significantly between the PBS and Ghrelin conditions. The predictor matrix was normalised by subtracting the predictor matrix by its mean and dividing by the L2 norm of the matrix, using the function RidgeCV. The β weights were computed analytically using the following formula:

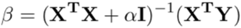

where X is the predictor matrix, α is the ridge penalty term, I is the identity matrix and Y is the observed dFz. Once fitted, the performance of the linear encoding model was assessed by using the independent test set to compute the explained variance (5-fold, cross-validated R2) value, or the coefficient of determination, defined as:

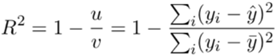

where u is residual sum of squares, v is the total sum of squares, yi is the photometry signal at index i, y^ is the predicted photometry signal at index i and y is the mean amplitude of the photometry signal in the test set. Linear models were estimated separately for data from individual animals.

### Electrophysiology

#### Slice preparation

Hippocampal recordings were studied in acute transverse slices as described previously (AlSubaie et al., 2021; Sánchez-Bellot et al., 2022; Wee and MacAskill, 2020). Mice were anaesthetized with a lethal dose of ketamine and xylazine, and perfused intracardially with ice-cold external solution containing (in mM): 190 sucrose, 25 glucose, 10 NaCl, 25 NaHCO3, 1.2 NaH2PO4, 2.5 KCl, 1 Na+ ascorbate, 2 Na+ pyruvate, 7 MgCl2 and 0.5 CaCl2, bubbled with 95% O2 and 5% CO2. Slices (400 μm thick) were cut in this solution and then transferred to artificial cerebrospinal fluid (aCSF) containing (in mM): 125 NaCl, 22.5 glucose, 25 NaHCO3, 1.25 NaH2PO4, 2.5 KCl, 1 Na+ ascorbate, 3 Na+ pyruvate, 1 MgCl2 and 2 CaCl2, bubbled with 95% O2 and 5% CO2. After 30 min at 35 °C, slices were stored for 30 min at 24 °C. All experiments were conducted at room temperature (22–24 °C). All chemicals were from Sigma, Hello Bio or Tocris.

Whole-cell recordings were performed on retrogradely labelled hippocampal pyramidal neurons with retrobeads visualised by their fluorescent cell bodies and targeted with Dodt contrast microscopy. For sequential paired recordings, neighbouring neurons were identified using a 40x objective at the same depth into the slice. The recording order of neuron pairs was alternated to avoid complications due to rundown. Borosilicate recording pipettes (3 – 5 M) were filled with different internal solutions depending on the experiment. For electrical stimulation experiments a Cs-gluconate based internal was used containing (in mM): 135 Gluconic acid, 10 HEPES, 7 KCl, 10 Na-phosphocreatine, 4 MgATP, 0.4 NaGTP, 10 TEA and 2 QX-314. Excitatory and inhibitory currents were electrically isolated by setting the holding potential at -70 mV (excitation) and 0 mV (inhibition) and recording in the presence of APV (50 μM). Alternatively, to record inhibitory miniature currents at -70 mV we used a high chloride internal (in mM): 135 CsCl, 10 HEPES, 7 KCl, 10 Na-phosphocreatine, 10 EGTA, 4 MgATP, 0.3 NaGTP, 10 TEA and 2 QX-314 in the presence of APV (50 μM), NBQX (10 μM) and TTX (1 μM) to block synaptic excitation and spontaneous IPSCs. Note for inhibitory mIPSC experiments (**Figure 5E-J**), electrophysiological recording was only possible for vS-NAc neurons but not vS-LH neurons in two ghrelin-injected animals as there was poor retrograde labelling of vS-LH neurons (i.e. vS-NAc, n = 37 neurons from 5 animals; vS-LH, n = 21 neurons from 3 animals). Recordings were made using a Multiclamp 700B amplifier, with electrical signals filtered at 4 kHz and sampled at 10 kHz.

### Statistical analyses

All statistics were calculated using the Python packages scipy, pingouin and statsmodels. Summary data are reported throughout the figures as boxplots, which show the median, 75th and 95th percentile as bar, box and whiskers respectively. Individual data points are also superimposed to aid visualisation of variance. Example physiology and imaging traces are represented as the mean +/-s.e.m across experiments. For the majority of analyses presented, normality of data distributions was determined by visual inspection of the data points. All data were analysed using statistical tests described in the statistical summary. Correction for multiple comparisons was conducted using the Benjamini-Hochberg method, unless otherwise stated. The alpha level was defined as 0.05. No power analysis was run to determine sample size a priori. The sample sizes chosen are similar to those used in previous publications. Throughout the figures and text, the * symbol represents p < 0.05, unless otherwise stated, and n.s. stands for not significant. Animals were randomly assigned to a virus cohort (e.g. ChR2 versus GFP), and where possible the experimenter was blinded to each mouse’s virus assignment when the experiment was performed. This was sometimes not possible due to e.g. the presence of the injection site in the recorded slice.

## ACKNOWLEDGEMENTS

We thank members of the MacAskill laboratory for helpful comments on the manuscript. A.F.M. was supported by a Sir Henry Dale Fellowship jointly funded by the Wellcome Trust and the Royal Society (grant number 109360/Z/15/Z) and by a UCL Excellence Fellowship. R.W.S.W. was supported by a UCL Graduate Research Scholarship and a UCL Overseas Research Scholarship. K.M. was supported by the Wellcome Trust 4-year PhD in Neuroscience at UCL (grant number 215165/Z/18/Z).

## STATSTICS SUMMARY

1 STATSTICS SUMMARY
| Figure | Descriptors | n | Test used | Statistic | p-value |
| --- | --- | --- | --- | --- | --- |
| 1A | Chow consumed PBS<br>vs. ghrelin | 22 mice | Paired t-test | $t_{(21)} = 9.85$ | $2.5 \times 10^{-9}$ |
| 1B | Frequency of investigation<br>Frequency of eating<br>Frequency of other, non<br>feeding behaviour | 22 mice | Paired t-test | $t_{(21)} = 0.35$<br>$t_{(21)} = 9.98$<br>$t_{(21)} = 3.23$ | 0.73<br>$2.0 \times 10^{-9}$<br>0.004 |
| 1C | P(inv->eat) | 22 mice | Paired t-test | $t_{(21)} = 10.94$ | $3.95 \times 10^{-10}$ |
| 2E | AUC Investigation<br>AUC Eating<br>AUC Other | 6 mice | Paired t-test (cf. baseline) | $t_{(5)} = 2.74$<br>$t_{(5)} = 4.67$<br>$t_{(5)} = 1.32$ | 0.04<br>0.01<br>0.24 |
| 2E | $\beta$ Investigation<br>$\beta$ Eating<br>$\beta$ Other | 6 mice | Paired t-test (cf. baseline) | $t_{(5)} = 2.58$<br>$t_{(5)} = 1.76$<br>$t_{(5)} = 1.01$ | 0.048<br>0.14<br>0.36 |
| 2H | AUC Investigation<br>AUC Eating<br>AUC Other | 6 mice | Paired t-test (cf. baseline) | $t_{(5)} = 1.38$<br>$t_{(5)} = 0.23$<br>$t_{(5)} = 0.98$ | 0.23<br>0.83<br>0.37 |
| 2H | $\beta$ Investigation<br>$\beta$ Eating<br>$\beta$ Other | 6 mice | Paired t-test (cf. baseline) | $t_{(5)} = 0.6$<br>$t_{(5)} = 0.57$<br>$t_{(5)} = 0.11$ | 0.56<br>0.59<br>0.91 |
| 3C | AUC Investigation<br>AUC eating<br>(PBS vs ghrelin) | 6 mice | Paired t-test | $t_{(5)} = 4.89$<br>$t_{(5)} = 1.57$ | 0.005<br>0.18 |
| 3D | $\beta$ Investigation<br>$\beta$ eating<br>(PBS vs ghrelin) | 6 mice | Paired t-test | $t_{(5)} = 2.92$<br>$t_{(5)} = 0.09$ | 0.03<br>0.93 |
| 3E | Food consumed x AUC<br>Food consumed x $\beta$ | 6 mice PBS + ghrelin | Linear regression | $R^2 = -0.72$<br>$R^2 = -0.70$ | 0.018<br>0.011 |
| 3H | AUC Investigation<br>AUC eating<br>(PBS vs ghrelin) | 6 mice | Paired t-test | $t_{(5)} = 0.68$<br>$t_{(5)} = 0.84$ | 0.52<br>0.44 |
| 3I | $\beta$ Investigation<br>$\beta$ eating<br>(PBS vs ghrelin) | 6 mice | Paired t-test | $t_{(5)} = 0.23$<br>$t_{(5)} = 1.01$ | 0.83<br>0.36 |
| 3J | Food consumed x AUC<br>Food consumed x $\beta$ | 6 mice PBS + ghrelin | Linear regression | $R^2 = -0.30$<br>$R^2 = -0.09$ | 0.35<br>0.77 |
| 4B | Investigation frequency<br>Eating Frequency | 6 mice | Paired t-test | $t_{(5)} = 1.57$<br>$t_{(5)} = 1.17$ | 0.18<br>0.30 |
| 4C | Investigation frequency<br>Eating Frequency | 5 mice | Paired t-test | $t_{(4)} = 0.38$<br>$t_{(4)} = 2.97$ | 0.72<br>0.04 |
| 4D | P(inv->eat) | 6 mice | Paired t-test | $t_{(4)} = 0.16$ | 0.88 |
| 4E | P(inv->eat) | 5 mice | Paired t-test | $t_{(4)} = 4.32$ | 0.01 |
| 5C | E:I ratio | 6 neurons (PBS)<br>9 neurons (ghrelin) | Independent t-test | $t_{(14)} = 3.98$ | 0.002 |
| 5D | E:I ratio | 6 neurons (PBS)<br>11 neurons (ghrelin) | Independent t-test | $t_{(16)} = 1.59$ | 0.13 |
| 5F | mIPSC amplitude<br>mIPSC frequency | 26 neurons (PBS)<br>37 neurons (ghrelin) | Independent t-test | $t_{(62)} = 2.79$<br>$t_{(62)} = 1.99$ | 0.007<br>0.06 |
| 5G | mIPSC amplitude<br>mIPSC frequency | 26 neurons (PBS)<br>21 neurons (ghrelin) | Independent t-test | $t_{(46)} = 1.20$<br>$t_{(46)} = 0.96$ | 0.23<br>0.34 |
| 6C | mIPSC amplitude<br>Scrambled<br>GHSR1a RNAi | 13 neurons (PBS)<br>14 neurons (ghrelin)<br>11 neurons (PBS)<br>17 neurons (ghrelin) | Independent t-test | $t_{(26)} = 4.17$<br>$t_{(27)} = 0.76$ | 0.0003<br>0.45 |
| 6F | AUC Investigation<br>Scrambled<br>GHSR1a RNAi | 8 mice<br>8 mice | Paired t-test | $t_{(7)} = 2.80$<br>$t_{(7)} = 0.75$ | 0.03<br>0.48 |
| 7B | Investigation frequency<br>Meal frequency | 6 mice (Scram)<br>6 mice (RNAi) | Independent t-test | $t_{(5)} = 0.84$<br>$t_{(5)} = 0.34$ | 0.42<br>0.73 |
| 7C | Investigation frequency<br>Meal frequency | 6 mice (Scram)<br>6 mice (RNAi) | Independent t-test | $t_{(5)} = 0.34$<br>$t_{(5)} = 2.44$ | 0.74<br>0.03 |
| 7D | P(inv->eat) | 6 mice | Independent t-test | $t_{(5)} = 0.45$ | 0.66 |
| 7E | P(inv->eat) | 6 mice | Independent t-test | $t_{(5)} = 4.69$ | 0.001 |

## SUPPLEMENTARY FIGURES

**Supplementary Figure 1.**
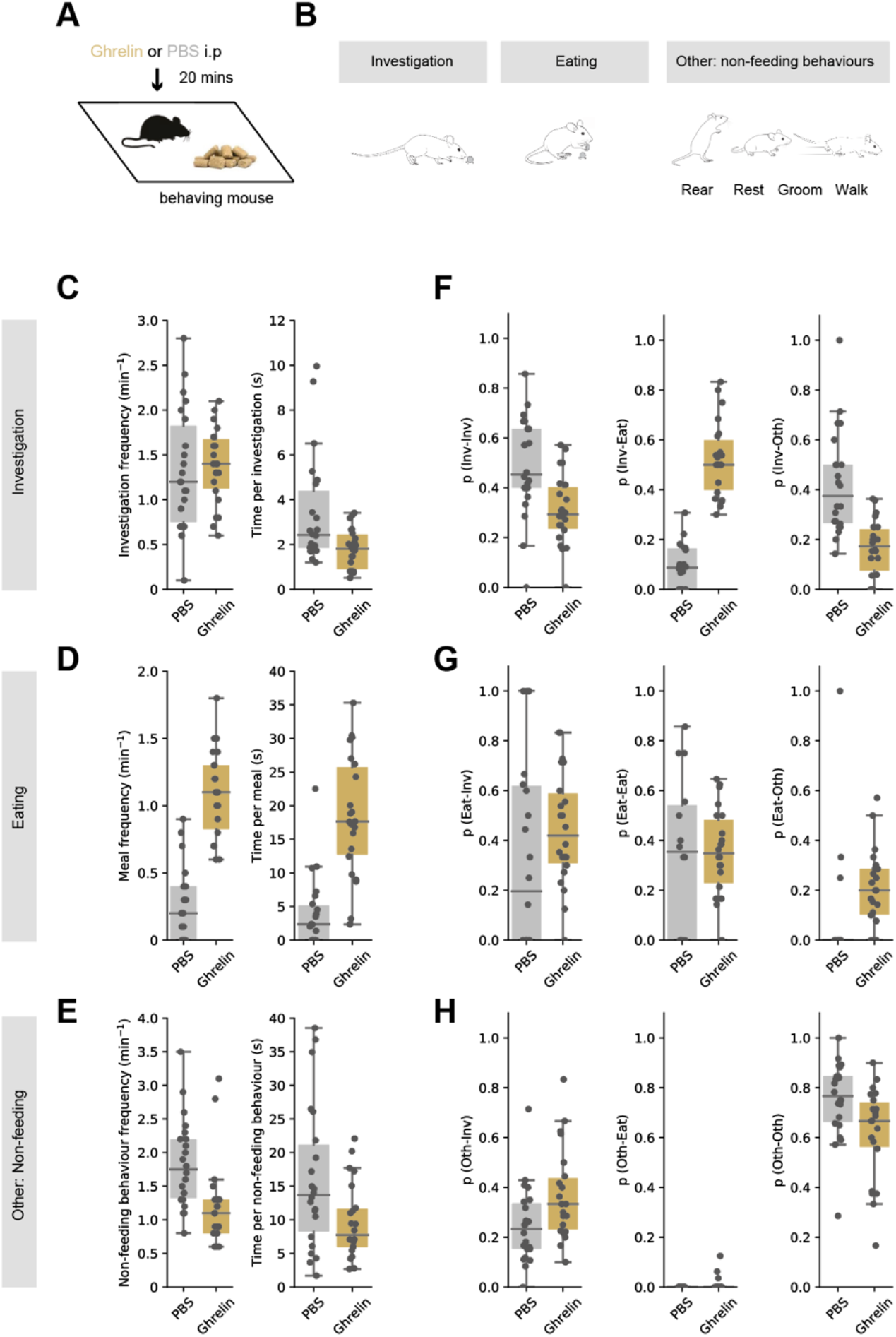
Expanded dataset for behaviour analysis in Figure 1. **A**. Top, schematic of experiment. Mouse was injected with either ghrelin or control PBS before exploring a well habituated chamber containing familiar chow for 10 min. Bottom, ghrelin administration (gold) increases chow consumption compared to PBS control (grey). **B**. Outline of how different behaviours were clustered for analysis. **C-E**. Model agnostic analysis of food investigation, eating, and non-feeding behaviours including grooming, quiet resting and rearing. Plots frequency of eat behaviour across the session (left) and duration of each bout (right). **F-H**. Markov analysis of feeding behaviour during ten-minute session as shown in main figure, but displayed as box plots with individual mice.

**Supplementary Figure 2.**
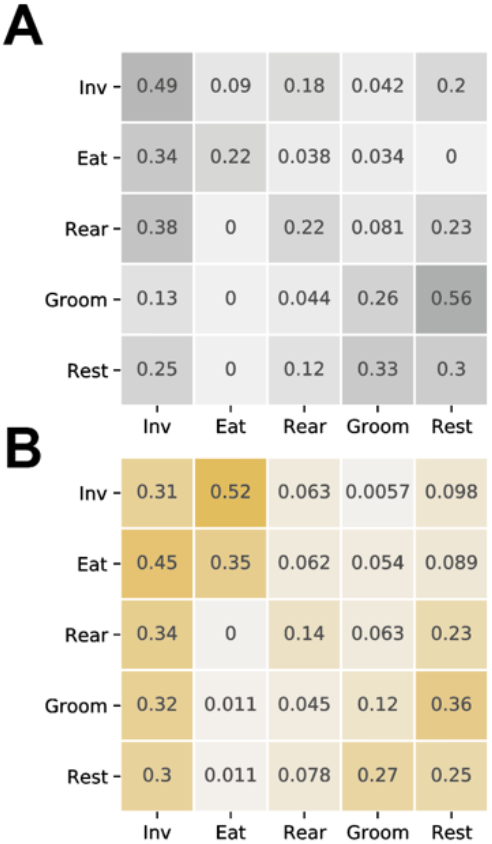
Expanded 5 × 5 Markov analysis of feeding behaviour during ten-minute session. State transition matrix for PBS (**A**) and ghrelin (**B**) treated mice. Each column and row add to one, and so the matrix represents the probability of the next behavioural state given the current state. Note that ghrelin increases the transition from investigation to eating, with minimal influence on other behavioural transitions. Note behaviours classed as ‘Oth’ in main analysis in **Figure 1** (Rear, Groom, Rest) do not undergo substantial change after ghrelin.

**Supplementary Figure 3.**
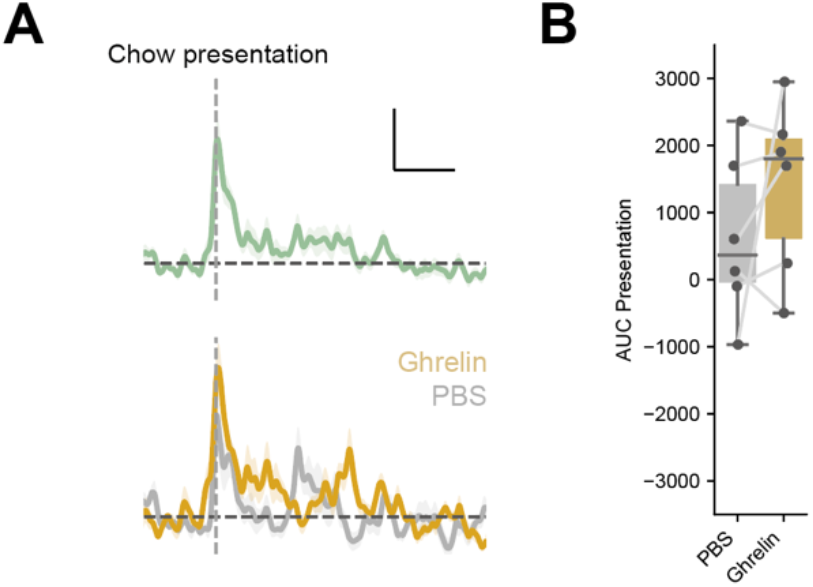
vS-LH neurons are active around chow presentation. **A**. Top, summary of activity around chow presentation for vS-LH neurons. B. Summary of activity around chow presentation after treatment with either PBS or ghrelin. **B**. Activity around presentation summarised as the area under the curve (AUC) of event-aligned activity. Note consistent increase in activity that is not influenced by peripheral ghrelin.

**Supplementary Figure 4.**
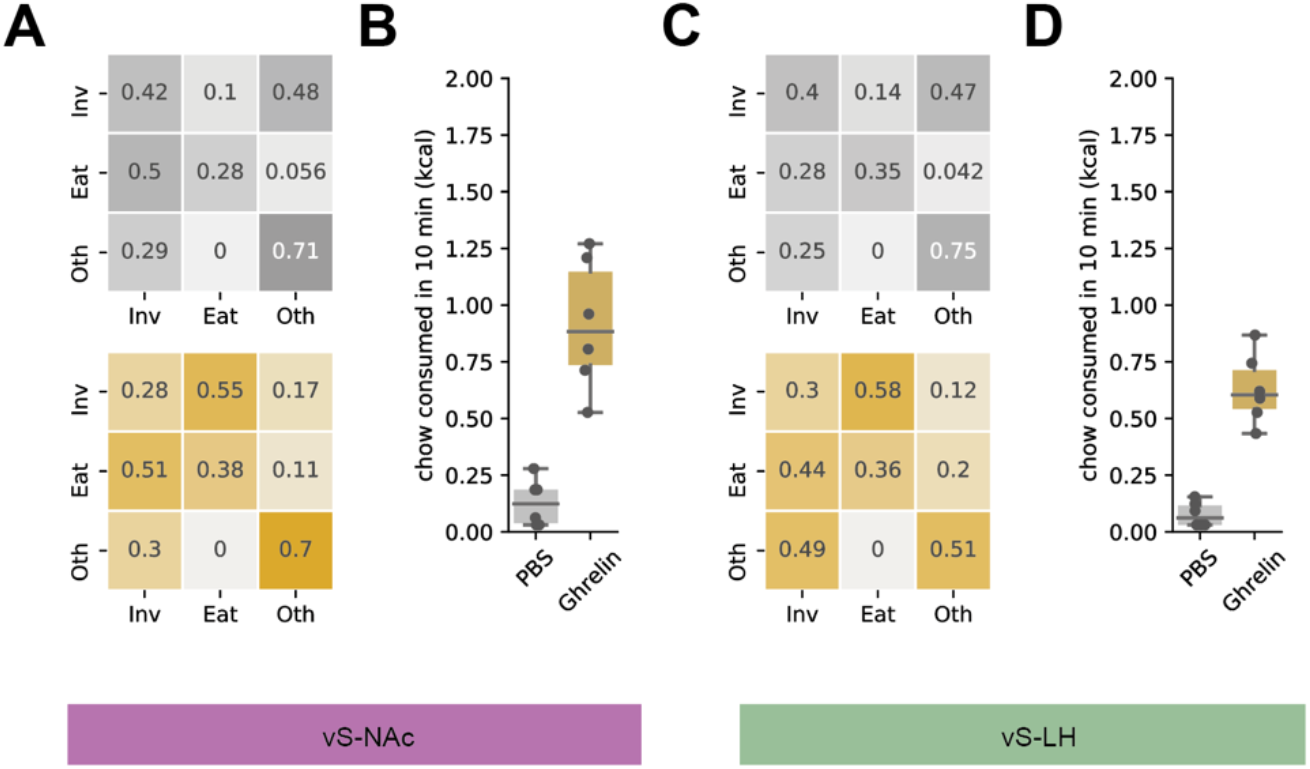
Behaviour of mice used for vS-NAc and vS-LH recordings is equivalent. **A**. Markov analysis of feeding behaviour of mice used for vS-NAc recordings during ten-minute session. Transition matrix for PBS (top) and ghrelin (bottom) treated mice. Each column and row add to one, and so the matrix represents the probability of the next behavioural state given the current state. Note that ghrelin increases the transition from investigation to eating, with minimal influence on other behavioural transitions. **B**. Ghrelin administration (gold) increases chow consumption compared to PBS control (grey). **C**,**D**. As in (**A**,**B**) but for vS-LH neurons.

**Supplementary Figure 5.**
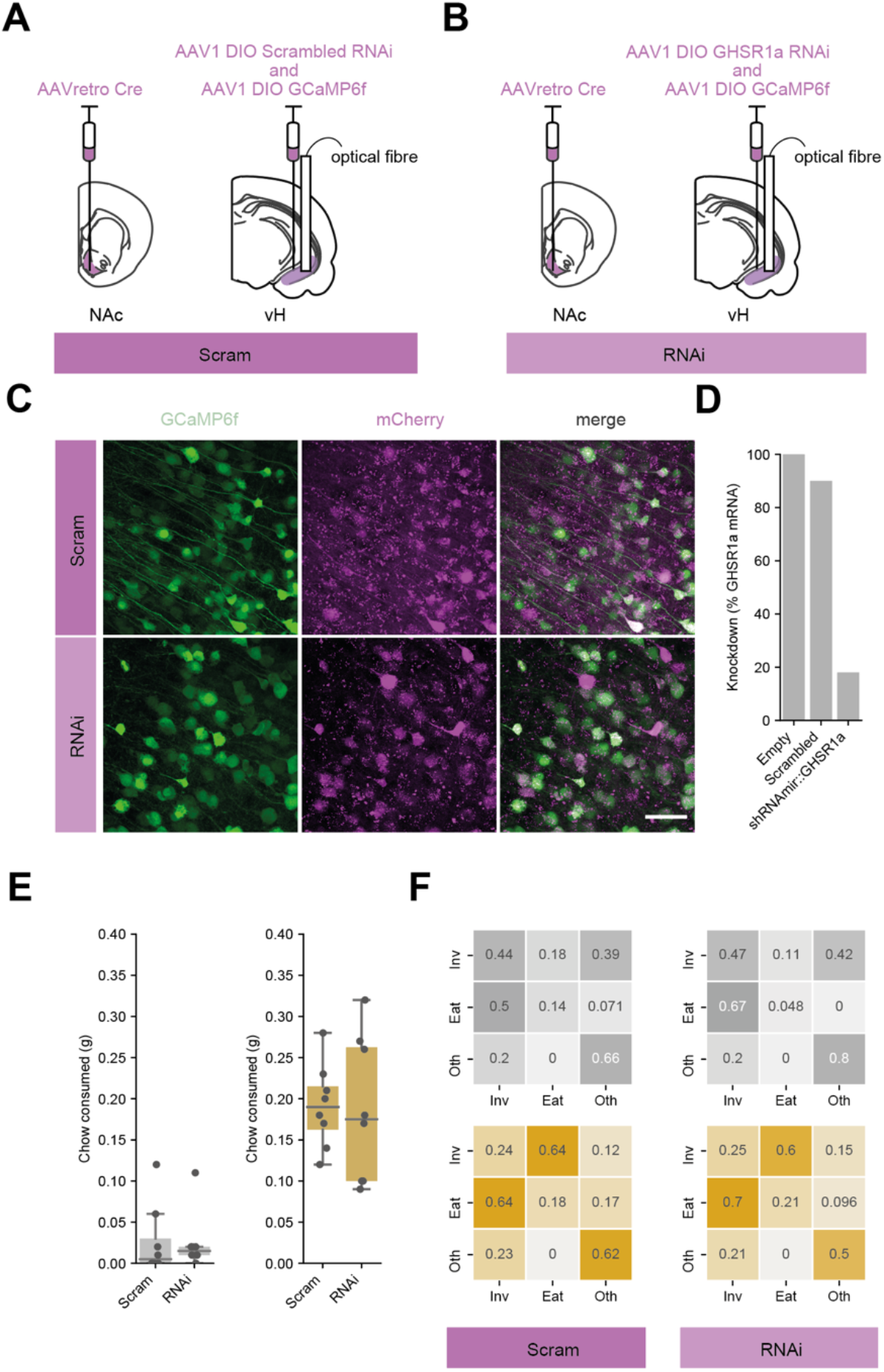
Characterisation of GHSR1a RNAi and demonstration that unilateral GHSR1a knockdown does not affect behaviour. **A, B**. Schematic of injections allowing intersectional targeting of vS-NAc neurons with either scrambled control RNAi (**A**), or GHSR1a RNAi (**B**) as well as GCaMP6f and an optical fibre to allow photometry recordings. **C**. Example images showing co-expression of GCaMP6f (green) with RNAi constructs (purple) in vS-NAc neurons after intersectional targeting. **D**. Quantification of GHSR1a levels in cell lines showing knockdown of GHSR1a mRNA after expression of GHSR1a RNAi but not scrambled RNAi compared to no vector control. **E**,**F**. Behavioural analysis showing no effect of unilateral GHSR1a RNAi expression of total chow consumed (**E**), or transition matrices (**F**). Note this lack of effect is in contrast to bilateral expression in **Figure 7**.

**Supplementary Figure 6.**
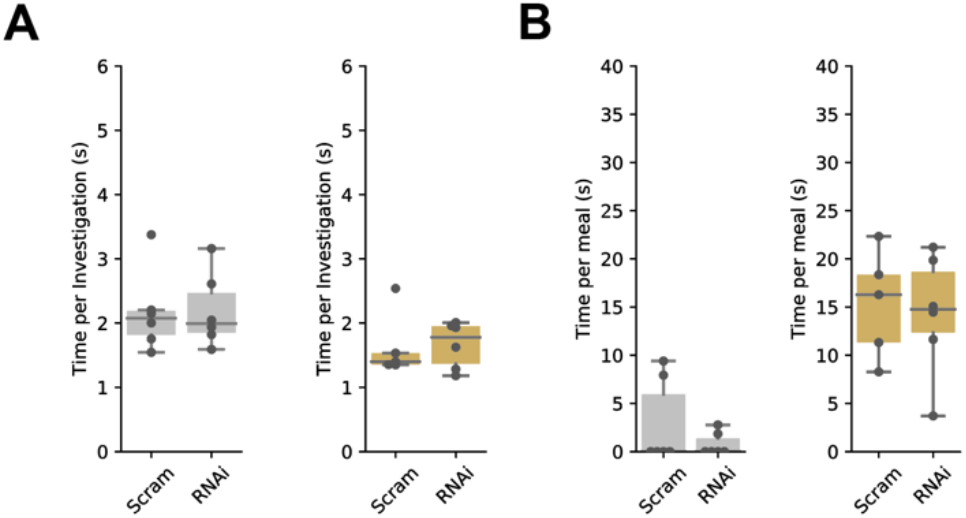
Bilateral GHSR1a knockdown in vS-NAc neurons has no effect on investigation or eating duration. **A**. Time spent per investigative bout in PBS (grey) and ghrelin (gold) treated mice expressing either scrambled RNAi or GHSR1a RNAi. **B**. As in (**A**) but time spent per eating bout. Note no differences across any conditions.

## SUPPLEMENTARY STATSTICS TABLE

| Figure | Descriptors | n | Test used | Statistic | p-value |
| --- | --- | --- | --- | --- | --- |
| S1C | Frequency of investigation<br>Time per Investigation | 22 mice | Paired t-test | $t_{(21)} = 0.35$<br>$t_{(21)} = 3.21$ | 0.73<br>0.025 |
| S1D | Frequency of eating<br>Time per eating | 22 mice | Paired t-test | $t_{(21)} = 9.98$<br>$t_{(21)} = 7.80$ | $1.19 \times 10^{-8}$<br>$7.43 \times 10^{-7}$ |
| S1E | Frequency of 'other'<br>Time per 'other' | 22 mice | Paired t-test | $t_{(21)} = 9.98$<br>$t_{(21)} = 2.61$ | 0.023<br>0.097 |
| S1F | P(inv->inv)<br>P(inv->eat)<br>P(inv->oth) | 22 mice | Paired t-test | $t_{(21)} = 4.74$<br>$t_{(21)} = 10.94$<br>$t_{(21)} = 4.99$ | 0.001<br>$3.56 \times 10^{-9}$<br>0.0006 |
| S1G | P(eat->inv)<br>P(eat->eat)<br>P(eat->oth) | 22 mice | Paired t-test | $t_{(21)} = 1.01$<br>$t_{(21)} = 1.45$<br>$t_{(21)} = 2.34$ | 1.0<br>1.0<br>0.27 |
| S1H | P(oth->inv)<br>P(oth->eat)<br>P(oth->oth) | 22 mice | Paired t-test | $t_{(21)} = 2.35$<br>$t_{(21)} = 1.60$<br>$t_{(21)} = 2.50$ | 0.26<br>1.0<br>0.19 |
| S3B | AUC Presentation<br>(PBS vs ghrelin) | 6 mice | Paired t-test | $t_{(5)} = 1.18$ | 0.29 |
| S4B,D | Chow consumed<br>PBS vs ghrelin (NAc)<br>PBS vs ghrelin (LH)<br>Nac vs LH (Ghrelin) | 6 mice | Paired t-test | $t_{(5)} = 5.36$<br>$t_{(5)} = 11.95$<br>$t_{(5)} = 2.12$ | 0.009<br>0.0002<br>0.18 |
| S5E | Chow consumed<br>Scram vs RNAi (PBS)<br>Scram vs RNAi (ghrelin) | 6 mice | Independent t-test | $t_{(5)} = 0.06$<br>$t_{(5)} = 0.13$ | 0.95<br>0.89 |
| S6A | Time per investigation<br>Scram vs RNAi (PBS)<br>Scram vs RNAi (ghrelin) | 6 mice | Independent t-test | $t_{(5)} = 0.06$<br>$t_{(5)} = 0.11$ | 0.95<br>0.91 |
| S6B | Time per eating<br>Scram vs RNAi (PBS)<br>Scram vs RNAi (ghrelin) | 6 mice | Independent t-test | $t_{(5)} = 1.11$<br>$t_{(5)} = 0.27$ | 0.29<br>0.79 |

